# Functional brain network correlates of pubertal timing and depressive symptoms in preadolescence

**DOI:** 10.64898/2026.05.22.727009

**Authors:** Athanasia Metoki, Benjamin P. Kay, Roselyne Chauvin, Samuel R. Krimmel, Anxu Wang, Philip N. Cho, Julia Monk, Noah J. Baden, Kristen M. Scheidter, Scott Marek, Timothy O. Laumann, Evan M. Gordon, Nico U. F. Dosenbach, Deanna M. Barch

**Author notes:** Address correspondence to Athanasia Metoki, PhD at or.

## Abstract

**BACKGROUND:** Variation in pubertal maturation relative to same-age, same-sex peers (pubertal timing) has been linked to increased risk for depressive symptoms during adolescence. This developmental period is also characterized by substantial reorganization of functional brain networks. However, how pubertal timing relates to resting-state functional connectivity (rsFC) changes and depression risk remains unclear.

**METHODS:** We examined pubertal timing and rsFC associations in preadolescents aged 9–11 years from the Adolescent Brain Cognitive Development (ABCD) Study. Pubertal timing was estimated using a puberty age gap approach based on parent-reported physical development. Linear mixed-effects and Bayesian multilevel models were used to assess cross-sectional and longitudinal associations between pubertal timing and rsFC across large-scale functional brain networks. We also tested whether rsFC differences explained associations between pubertal timing and later depressive symptoms.

**RESULTS:** Earlier pubertal timing was associated with heterogeneous rsFC patterns, with stronger and more widespread effects in females. In females, earlier pubertal timing was associated with rsFC increases and decreases across sensory–motor and association networks, whereas in males, associations were more limited and localized to sensorimotor and cerebellar systems. Longitudinally, earlier pubertal timing in females predicted reductions in rsFC at the 2-year follow-up, with no significant associations in males. rsFC differences did not explain the pubertal timing and later depressive symptoms association.

**CONCLUSIONS:** Pubertal timing is associated with sex-specific patterns of brain functional connectivity during early adolescence, with greater heterogeneity and broader network involvement in females. These findings suggest that pubertal maturation contributes to early reorganization of functional brain networks, although these changes did not explain subsequent depressive symptoms.

## Introduction

Adolescence marks a period of profound biological, psychological, and social transition, during which pubertal development reshapes multiple aspects of brain and behavior (1–6). During this time, large-scale functional brain networks undergo substantial reorganization, reflected in changes in resting-state functional connectivity (rsFC) within and between distributed neural systems (7–11). However, adolescents vary considerably in the timing of pubertal maturation relative to their same-sex peers, a construct referred to as pubertal timing. These individual differences represent an important source of variability in neurodevelopment and may influence functional brain network organization during adolescence.

Understanding how variation in pubertal timing relates to the development of large-scale functional brain networks may provide insight into emerging mental health risk during this period. One outcome consistently linked to pubertal timing is depression. Depressive symptoms increase sharply during early adolescence (12–14), with girls showing faster and larger increases than boys (12,15–17). Earlier pubertal timing has been associated with more persistent depressive symptoms, earlier onset of internalizing disorders, and greater likelihood of clinical diagnoses (18–27). These effects are typically stronger in girls, whereas findings in boys are more mixed and remain comparatively understudied (24,28–38). Despite growing evidence linking pubertal timing to both brain development and mental health, how variation in pubertal timing shapes large-scale functional brain network organization and relates to depressive symptoms remains poorly understood.

Functional neuroimaging studies have begun to examine associations among pubertal timing, brain connectivity, and depressive symptoms, but existing findings remain mixed. Vijayakumar et al. (39) using combined data from baseline and 2-year follow-up assessments from the Adolescent Brain Cognitive Development® (ABCD; (40)), reported that earlier pubertal timing was associated with reduced resting-state functional connectivity (rsFC) between limbic regions (amygdala, hippocampus) and cortical networks (i.e., the cingulo-opercular, sensorimotor mouth, and ventral attention). These rsFC reductions mediated the association between earlier pubertal timing and greater depressive symptoms. In contrast, Colich et al. (41) found that earlier pubertal timing was associated with higher externalizing, but not internalizing, symptoms in both boys and girls and that amygdala–mPFC connectivity showed no association with pubertal timing or psychopathology. Lastly, in a very small, preliminary study of 68 adolescents from an independent sample, Chahal et al. (42) reported that earlier pubertal timing was associated with reduced connectivity among regions involved in affective and self-referential processing (i.e., cingulate gyrus, precuneus, insula, inferior parietal lobule) with these patterns associated with concurrent and later depressive symptoms. Together, these findings suggest that pubertal timing may relate to both rsFC and depressive symptoms, although existing findings remain inconsistent and are largely limited to selected network subsets.

A key limitation of prior work is the reliance on a priori–defined circuits, which may overlook broader network-level effects. Because results are constrained to a limited set of predefined pathways, reported associations may vary depending on the specific circuits and modeling approaches examined. In contrast, data-driven, connectome-wide analyses provide a more comprehensive assessment of brain organization but require large samples as brain-behavior effect sizes are often small and variable across connections (43). Large-scale datasets therefore offer an opportunity to identify more reliable and generalizable associations between pubertal timing, brain connectivity, and mental health.

Here, we leverage the large ABCD cohort (n = 11,875) to conduct a data-driven, whole-connectome analysis of pubertal timing, brain connectivity, and depressive symptoms in early adolescence. Using resting-state fMRI from youth assessed at baseline (BL; ages 9–11) and the 2-year follow-up (Y2; ages 11–13), we quantified rsFC across 171 network-pair connections spanning major cortical networks and subcortical structures. Pubertal timing was estimated at BL using a puberty age gap approach derived from parent-reported physical development, and depressive symptoms were assessed at the BL, Y2, and the 3-year follow-up (Y3; ages 12–14) using the CBCL Withdrawn/Depressed subscale. First, we examined whether pubertal timing was associated with rsFC at BL (rsFC_BL_), and whether the strength and direction of these associations varied across large-scale network pairs. Next, we tested whether pubertal timing predicted longitudinal changes in rsFC. We then evaluated whether pubertal timing was associated with depressive symptoms both concurrently and prospectively, establishing the relevance of pubertal timing to mental health outcomes in this cohort. Finally, we assessed whether longitudinal differences in rsFC mediated the relationship between pubertal timing and later depressive symptoms.

## Methods and Materials

### Participants

Data for the present study were drawn from the ABCD Study curated annual release 5.1. The analytic sample for the present study included multiple nested samples depending on data availability across behavioral and neuroimaging modalities (Fig. 1A). Details on sample construction and demographic characteristics are provided in the Supplemental Information (Fig. S1; Table S1).

**Figure 1.**
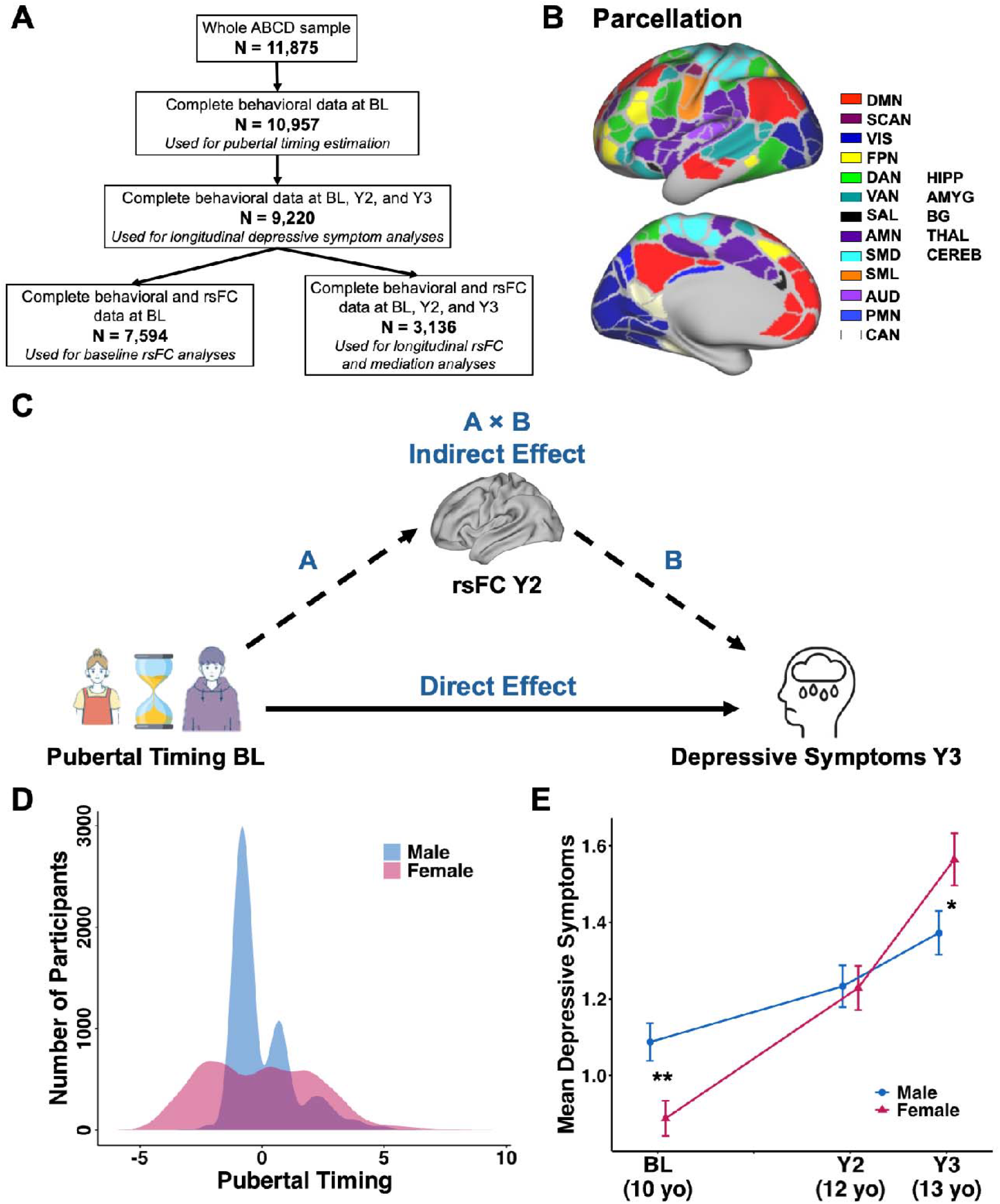
Sample derivation, parcellation, and sex differences in pubertal timing and depressive symptoms. (A) Flow diagram illustrating sample derivation. The full baseline (BL) cohort (n = 11,875) was sequentially restricted to participants with complete behavioral data at BL, complete behavioral data across BL and follow-ups, and complete resting-state functional connectivity (rsFC) data at BL and longitudinally. Subsamples used for pubertal timing calculation, depressive symptom analyses, cross-sectional rsFC analyses, and longitudinal rsFC and mediation models are indicated. (B) Parcellation. A region-of-interest (ROI) approach was employed, incorporating a total of 394 parcels: 333 cortical parcels defined by Gordon et al. (45) and 61 subcortical parcels from Seitzman et al. (46). The ROIs were averaged according to their functional network or anatomical grouping (i.e., anatomical structure) to generate timecourses for higher-order regions. The cortical ROIs were organized into canonical large-scale functional networks: default mode (DMN), somato-cognitive action (SCAN), visual (VIS), frontoparietal (FPN), dorsal attention (DAN), ventral attention (VAN), salience (SAL), action mode (AMN), sensorimotor dorsal (SMD), somatomotor lateral (SML), auditory (AUD), parietal memory (PMN), and context association (CAN). The subcortical ROIs were grouped into the following anatomical structures: hippocampus (HIPP), amygdala (AMYG), basal ganglia (BG), thalamus (THAL), and cerebellum (CEREB). (C) Longitudinal mediation model linking pubertal timing, rsFC, and depressive symptoms. Path A modeled associations between pubertal timing at BL and rsFC at the 2-year follow-up (Y2), whereas Path B modeled associations between rsFC at Y2 and depressive symptoms at the 3-year follow-up (Y3). Indirect effects (A × B) were estimated using Bayesian multilevel models. Analyses adjusted for baseline rsFC, prior depressive symptoms, and age at each time point. (D) Distribution of pubertal timing scores in males and females. Pubertal timing was operationalized as the difference between age predicted from physical development and chronological age. Positive values reflect earlier pubertal timing relative to same-sex, same-age peers, whereas negative values reflect later pubertal timing. Density plots illustrate sex differences in distributional shape, with females exhibiting a broader and more heterogeneous distribution and more extreme early and late values. The distribution of pubertal timing scores differed significantly between sexes (Wilcoxon rank-sum test, p < 0.001; Table S1). (E) Depressive symptom scores (CBCL Withdrawn/Depressed scale) at BL, Y2, and Y3 stratified by sex. Sex differences varied across time, with higher depressive symptoms in males at BL, no significant difference at Y2, and higher depressive symptoms in females at Y3 (Table S1). ** p < 0.001, * p < 0.01

### fMRI Data Acquisition and Processing

Resting-state fMRI data were obtained from the ABCD Study’s BL and Y2 assessments and were preprocessed as previously described (44). Once processed, rsFC timecourses were extracted from 333 cortical (45) and 61 subcortical (46) parcels. These regions of interest were then averaged according to their functional network (47,48) or anatomical grouping (i.e., anatomical structures) to generate higher-order regions’ timecourses (Fig. 1B). Pearson correlation coefficients were computed between all network/structure pair timecourses, yielding 171 unique connections per participant, and were normalized using Fisher’s r-to-z transformation (47). Detailed preprocessing steps, parcel definitions, and network assignments are provided in the Supplemental Information.

### Developmental and Behavioral Measures

#### Pubertal Timing

Pubertal timing was estimated separately for males and females using ABCD BL data and the puberty age method described by Dehestani et al. (48). Chronological age was predicted using items from the parent-reported Pubertal Development Scale (PDS) (49), including general physical development (e.g., height growth, body hair, skin changes) and sex-specific indicators (e.g., facial hair, voice deepening, breast development, menstruation). Parent reports were used rather than youth self-reports given evidence that younger participants tend to overestimate development (50). The difference between predicted age (puberty age) and the actual chronological age, adjusted for regression-to-the-mean bias, yielded the puberty age gap. Positive values indicate earlier pubertal timing relative to chronological age, while negative values indicate later timing. Details about the pubertal timing estimation procedure are provided in the Supplemental Information.

#### Depressive Symptoms

Depressive symptoms were assessed using the parent-reported CBCL (51) from ABCD’s BL, Y2, and Y3 assessments. Analyses used raw scores (52) from the Withdrawn/Depressed syndrome scale, which captures symptoms such as social withdrawal, lack of enjoyment, low energy, and feelings of sadness or depression. Due to the positively skewed and overdispersed distribution of depressive symptom scores, they were modeled using a negative binomial distribution with a log link in all relevant analyses.

#### Covariates

In all analyses, age at the assessment wave corresponding to the outcome variable was included as a covariate to account for developmental differences in pubertal timing, brain function, and depressive symptoms. In longitudinal analyses with depressive symptoms as the outcome, models additionally adjusted for prior depressive symptoms (e.g., Y2 for Y3 outcome). Models included random intercepts of site, family, and participant, with network-pair–specific random slopes in rsFC models, allowing pubertal timing–rsFC associations to vary across connections within a partially pooled hierarchical framework. Random intercepts for site and family accounted for non-independence due to shared site and family membership, whereas the participant-level random intercept accounted for repeated measurements within individuals across network pairs.

### Statistical Analyses

#### Baseline associations between pubertal timing and rsFC

To characterize how pubertal timing relates to large-scale functional brain organization at BL, we examined associations between pubertal timing and rsFC across network pairs using complementary frequentist and Bayesian multilevel approaches. We first fit linear mixed-effects models including a pubertal timing × sex interaction and network-pair–specific random slopes.

Sex-specific heterogeneity was evaluated by comparing pooled and extended models allowing female-specific deviations. Sex-stratified models were then fit to aid interpretation. To obtain network-pair–specific estimates, we fit Bayesian multilevel models with the same model structure as in the frequentist approach. Posterior estimates were used to characterize the magnitude, direction, and uncertainty of pubertal timing–rsFC associations across network pairs. Details of model specification, estimation procedures, model comparison, and Bayesian inference are provided in the Supplemental Information.

#### Longitudinal pubertal timing relations to rsFC

Next, we examined whether pubertal timing was associated with longitudinal differences in rsFC. Within each sex, rsFC at Y2 (rsFC_Y2_) was modeled as a function of pubertal timing, adjusting for age at Y2 (age_Y2_) and rsFC_BL_. Linear mixed-effects models included network-pair– specific random slopes for pubertal timing. To identify stable longitudinal associations for mediation analyses, we implemented a 5-fold cross-validation procedure using Bayesian multilevel models. Network pairs showing consistent effects across folds (≥3 of 5 training folds) were carried forward for mediation analyses, and final estimates were obtained in the full sample. Details of model specification, cross-validation procedures, and stability criteria are provided in the Supplemental Information.

#### Associations between pubertal timing and depressive symptoms, and network-specific mediation

To evaluate associations between pubertal timing and depressive symptoms, we conducted cross-sectional and longitudinal mixed-effects analyses. First, we examined whether pubertal timing was associated with depression_BL_ using linear mixed-effects models including pubertal timing, sex, and their interaction as predictors, adjusting for age at BL (age_BL_). We then tested whether pubertal timing predicted later depressive symptoms using longitudinal models with depression_Y3_ as the outcome, adjusting for depression_Y2_ and age at Y3 (age_Y3_). Together, these analyses assessed concurrent and prospective associations. Finally, we tested a longitudinal, network-specific mediation model using Bayesian multilevel analysis, linking pubertal timing to rsFC_Y2_ (Path A) and rsFC_Y2_ to depression_Y3_ (Path B), adjusting for rsFC_BL_ and prior depression_Y2_ (Fig. 1C). Details of model specification, Bayesian estimation, and mediation procedures are provided in the Supplemental Information.

## Results

### Youths’ demographic characteristics

In this sample (*n* = 9,220), males (52.5%) and females (47.5%) did not differ significantly in age_BL_. Pubertal timing was defined as age-adjusted pubertal status (positive values = earlier maturation; negative values = later maturation relative to same-age, same-sex peers). Because pubertal timing distributions differed between males and females and violated normality and homogeneity of variance assumptions, sex comparisons were conducted using nonparametric Wilcoxon rank-sum tests (Table S1). Females exhibited a broader, more heterogeneous distribution of pubertal timing, with more extreme early and late values, resulting in a significant sex difference (Fig. 1D; Table S1). Sex differences in depressive symptoms varied across time. Wilcoxon rank-sum tests indicated significantly higher depression_BL_ in males relative to females, no significant sex difference at Y2, and a reversal of this pattern by Y3, with females exhibiting higher depressive symptom scores than males (Fig. 1E; Table S1).

### Earlier pubertal timing shows sex-differentiated rsFC patterns

In the pooled model including both sexes, there was no evidence for an average association between pubertal timing and rsFC_BL_ or for a pubertal timing × sex interaction at the population level, indicating that their overall association did not differ by sex. Despite the absence of a population-level pubertal timing effect, there was clear evidence of heterogeneity, with pubertal timing–rsFC associations varying in both strength and direction across network pairs (Table S2). To assess whether this heterogeneity differed by sex, we compared the pooled model to an extended pooled model allowing network-pair–specific deviations in females relative to males. The extended model provided a significantly better fit, with more pronounced network-specific differences in pubertal timing–rsFC_BL_ associations in females (Table S3). Together, these results indicate substantial heterogeneity in pubertal timing–rsFC_BL_ associations across network pairs, with sex-specific differences in the magnitude of this variability.

Given the observed sex-specific network-level heterogeneity in the extended pooled model, sex-stratified mixed-effects models were fit to estimate associations between pubertal timing and rsFC_BL_ within each sex (Equation S1). In females, there was no evidence for an overall fixed-effect association between pubertal timing and rsFC_BL_, but network-pair–specific variability remained evident (Table S2). Bayesian multilevel models further revealed a heterogeneous pattern of associations across the connectome, with both negative and positive pubertal timing effects across 85 distinct network pairs (Fig. 2A). The strongest negative pubertal timing–rsFC_BL_ associations were observed for the basal ganglia (BG)–auditory (AUD) and BG–somatomotor lateral (SML) connections, indicating that earlier-maturing females showed lower rsFC_BL_ in these connections relative to their same-age, same-sex peers. The strongest negative effects (i.e., networks/anatomical structures with the largest median absolute negative effect sizes across all pairs involving that network/anatomical structure) were observed in BG and ventral attention (VAN), followed by the cerebellum (CEREB) and sensorimotor dorsal (SMD) (Fig. 3A). Networks most frequently involved in negative pubertal timing–rsFC_BL_ associations (based on the number of significant network pairs) were salience (SAL) and SML, followed by AUD, BG, and default mode (DMN). The strongest positive pubertal timing–rsFC_BL_ associations were observed for the somato-cognitive action (SCAN)–SMD and SML–SMD connections. At the network level, the strongest positive effects were found for the VIS and VAN, followed by SMD and parietal memory (PMN) (Fig. 3B). Networks most frequently involved in positive pubertal timing–rsFC_BL_ associations were CEREB, SMD, FPN, and dorsal attention (DAN).

**Figure 2.**
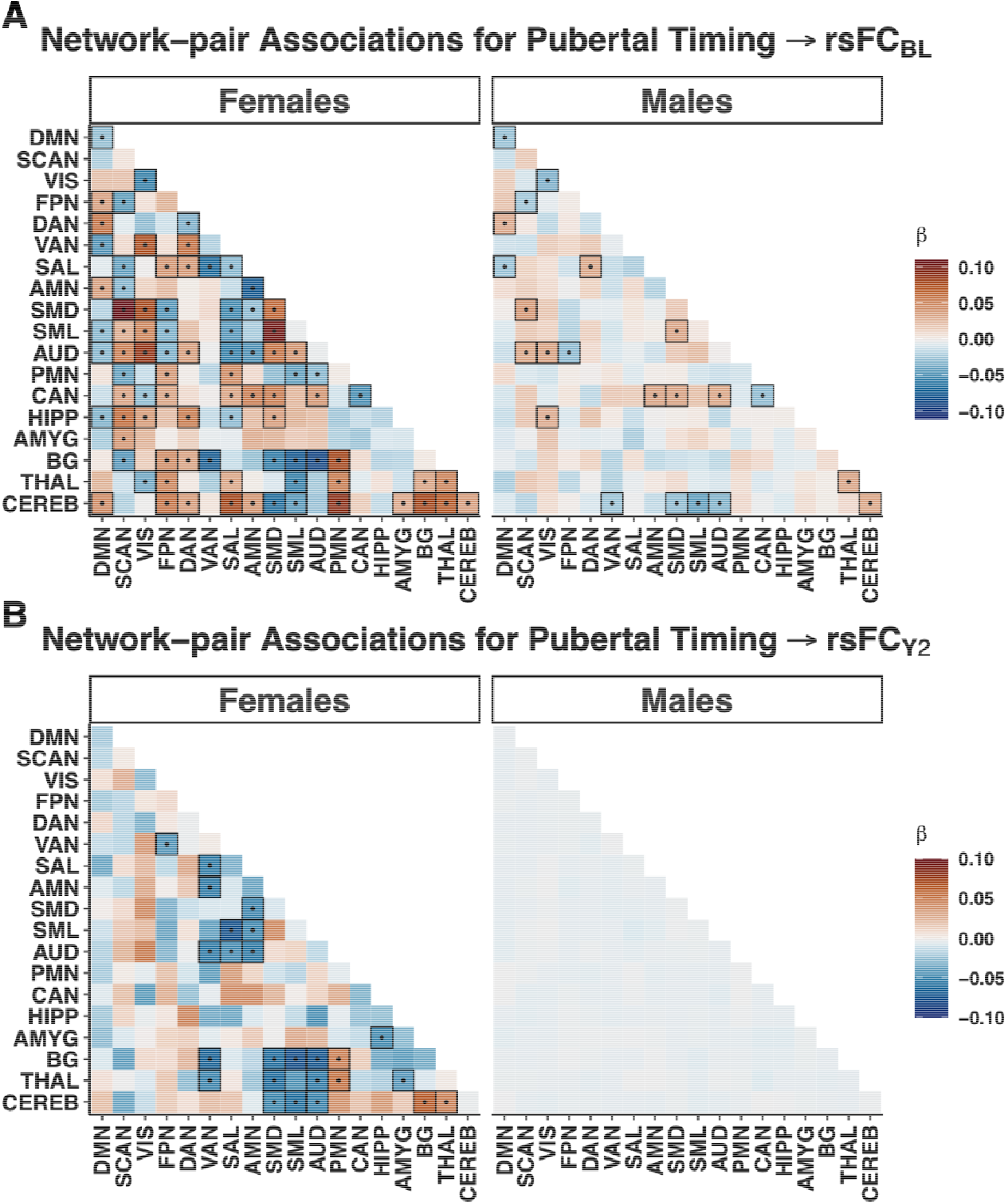
Network-pair associations between pubertal timing and resting-state functional connectivity at (A) baseline and (B) the 2-year follow-up. Heatmaps depict posterior median slopes (β) from Bayesian multilevel models estimating cross-sectional associations between pubertal timing and rsFC across large-scale functional network pairs. Warmer colors indicate positive associations (earlier pubertal timing associated with higher rsFC), whereas cooler colors indicate negative associations (earlier pubertal timing associated with lower rsFC). Black outlines and overlaid dots indicate network pairs whose 95% credible intervals excluded zero, reflecting credible cross-sectional associations under the hierarchical model (Panel A) and network pairs that met the predefined stability criterion (credible non-zero effect in ≥3 of 5 cross-validation folds with consistent direction), indicating reproducible longitudinal effects (Panel B).

**Figure 3.**
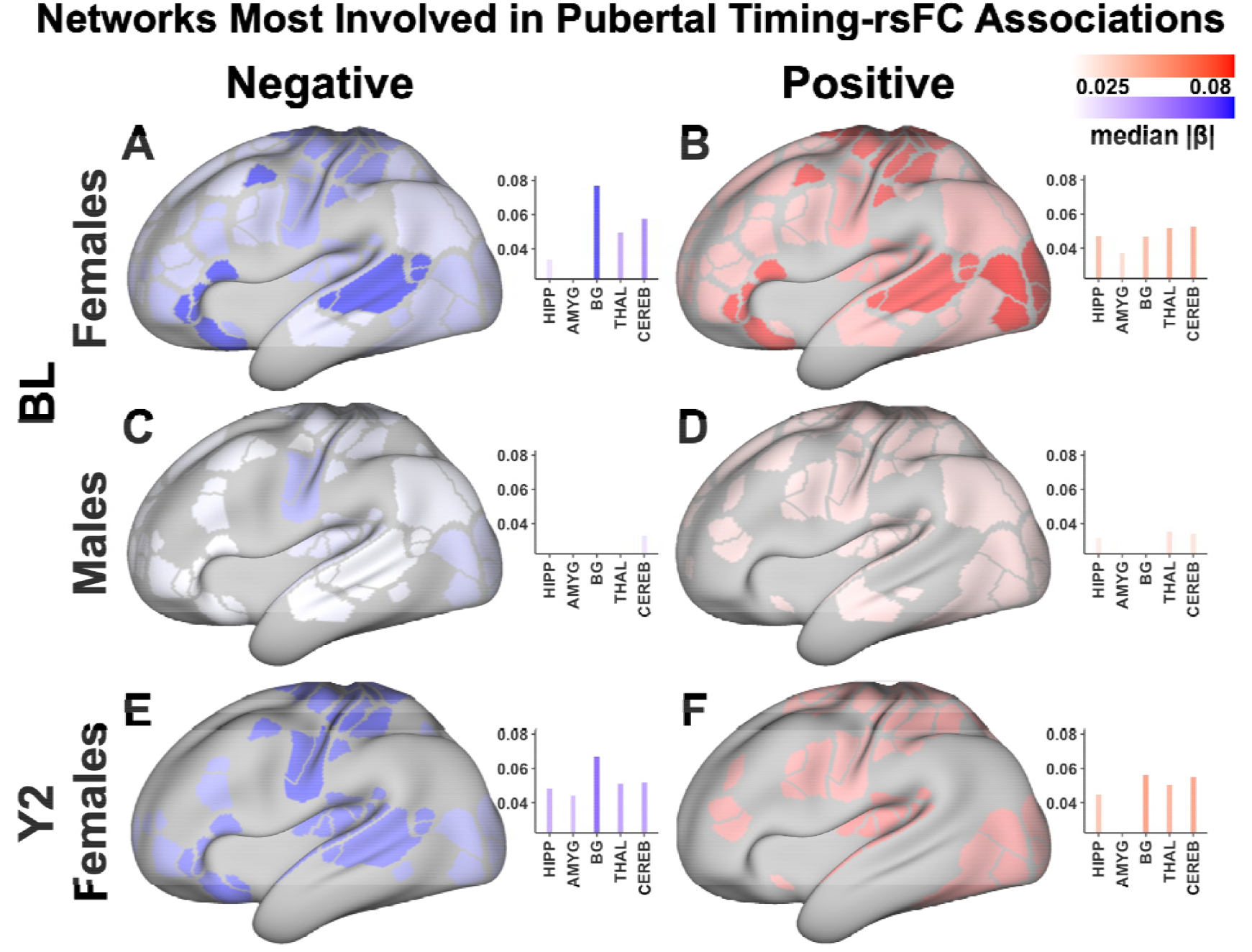
Network-level magnitude of positive and negative pubertal timing effects on resting-state connectivity across sex and time. Surface maps depict the median absolute effect size (|β|) of pubertal timing on resting-state functional connectivity (rsFC), separated by direction of association. Positive effects indicate higher rsFC whereas negative effects indicate lower rsFC in earlier-maturing youth. Left column shows networks most frequently involved in positive associations; right column shows networks most frequently involved in negative associations. Rows correspond to males at baseline (BL; top), females at BL (middle), and females at the 2-year follow-up (Y2; bottom). Color intensity reflects the median absolute β within each network, with white indicating smaller effects and darker red (positive column) or blue (negative column) indicating larger effect magnitudes. All panels use a common color scale (0.025–0.08) to allow direct comparison across sex and time.

In males, pubertal timing was likewise not associated with rsFC_BL_ at the overall (fixed-effect) level. Network-pair–specific variability in pubertal timing effects was smaller in magnitude compared to females, indicating less heterogeneity in pubertal timing–rsFC_BL_ associations across network pairs (Table S2). Bayesian multilevel models revealed pubertal timing–rsFC_BL_ associations across 22 distinct network pairs, reflecting a more limited pattern of both negative and positive effects (Fig. 2A; Fig. S2) compared to females. The strongest negative associations involved SML–CEREB and within-network visual (VIS) connectivity (VIS–VIS). At the network level, the strongest negative effects were observed in SML and VIS, followed by the AUD and CEREB (Fig. 3C). Networks most frequently involved in negative associations were CEREB and AUD, followed by DMN and FPN. In contrast, the strongest positive associations were observed for SCAN–SMD and VIS–AUD. The strongest positive effects involved SCAN and SMD, followed by context association (CAN) and action mode (AMN) (Fig. 3D). Networks most frequently involved in positive associations were CEREB, SMD, FPN, and DAN. Full network-pair–specific results for females and males are reported in Tables S4 and S5.

### Earlier pubertal timing predicts increased depressive symptoms over time

The hypothesis that rsFC mediates the effect of pubertal timing on depressive symptoms requires that pubertal timing be significantly associated with depressive symptoms over time. At BL (Equation S3), earlier pubertal timing was significantly associated with higher depression, indicating that youth who were more advanced in pubertal development compared to peers reported greater depressive symptom severity. The pubertal timing × sex interaction was not significant, indicating similar associations in males and females (Table S6). In longitudinal analyses (Equation S4) earlier pubertal timing predicted higher depression_Y3_ after adjusting for prior symptoms (depression_Y2_; Table S6). As expected, depressive symptoms showed strong temporal stability, with depression_Y2_ robustly predicting depression_Y3_. The pubertal timing × sex interaction was again not significant. Together, these findings indicate that pubertal timing is associated with depressive symptoms both concurrently and over time, supporting further examination of potential neural mechanisms linking the two.

### Female-specific longitudinal rsFC alterations are associated with pubertal timing, but do not mediate relations to depression

We next examined whether pubertal timing was associated with longitudinal changes in rsFC from BL to Y2 while adjusting for age_Y2_ and rsFC_BL_ (Equation S2). In females, earlier pubertal timing was associated with lower rsFC_Y2_ at the fixed-effect (population) level after accounting for rsFC_BL_ (Table S7). As expected, rsFC showed strong temporal stability, with rsFC_BL_ robustly predicting rsFC_Y2_. Importantly, there was heterogeneity in pubertal timing effects across network pairs, reflected in variability of network-pair–specific random slopes for pubertal timing (SD = 0.036; Table S7), indicating that longitudinal pubertal timing–rsFC_Y2_ associations varied meaningfully across large-scale functional network pairs.

To identify network pairs most consistently contributing to this heterogeneity, we performed a Bayesian 5-fold cross-validation analysis in females, estimating pubertal timing effects on rsFC_Y2_ while adjusting for age_Y2_ and rsFC_BL_. Network pairs were considered stable if 95% credible intervals excluded zero in ≥3 folds and the direction of the pubertal timing effect (positive or negative) was consistent across folds. This analysis identified a subset of 25 network pairs exhibiting reproducible longitudinal associations with pubertal timing across folds (Fig. 2B; Fig. S3). The strongest negative pubertal timing–rsFC_Y2_ associations were observed for the BG–SML and SAL–SML. At the network level, the strongest negative effects were found in BG and SML, followed by SMD and AUD (Fig. 3E). Networks most frequently involved in negative pubertal timing–rsFC_Y2_ associations were VAN and AUD, followed by BG, SML, and THAL. The strongest positive associations were BG–CEREB and PMN–THAL and the strongest positive effects were found in BG and CEREB (Fig. 3F). The structure most frequently involved in positive pubertal timing–rsFC_Y2_ associations was the BG.

In mediation models, pubertal timing showed a robust positive association with depression_Y3_, which remained after accounting for rsFC_Y2_, with a similarly positive direct effect, indicating that earlier pubertal timing was associated with higher later depressive symptoms independent of rsFC differences. While Path A (pubertal timing ➔ rsFC_Y2_, controlling for rsFC_BL_) showed effects with 95% credible intervals excluding zero for the 25 selected network pairs, Path B (rsFC_Y2_➔ depression_Y3_, controlling for depression_Y2_) showed no evidence of associations. Accordingly, no network-specific indirect effects showed credible evidence of mediation. Full cross-validation and mediation results for females are reported in Tables S8 and S9.

In males, pubertal timing was not significantly associated with rsFC_Y2_. In contrast to females, there was little evidence of heterogeneity in pubertal timing–rsFC_Y2_ associations across network pairs (Table S7). For consistency with the female analyses, we applied the same Bayesian 5-fold cross-validation procedure to identify rsFC_Y2_ network pairs showing stable longitudinal associations with pubertal timing. As expected, given the minimal between-network variability, no network pairs met the predefined stability criteria (Fig. 2B).

## Discussion

The current study investigated whether earlier pubertal timing is associated with large-scale patterns of rsFC in preadolescence and whether these functional network-level differences contribute to subsequent depressive symptoms. Leveraging a large, population-based cohort (ABCD Study) and connectome-wide multilevel modeling, we examined both cross-sectional and longitudinal associations across all major resting-state functional network connections. We found no evidence for a global association between pubertal timing and rsFC. Instead, earlier pubertal timing was linked to heterogeneous, network-specific, and sex-specific differences in connectivity. In addition, earlier pubertal timing was associated with higher concurrent and subsequent depressive symptoms, consistent with prior findings linking earlier pubertal maturation to increased depression risk during adolescence (18–22,29,30,37–39,41). However, rsFC did not mediate the association between pubertal timing and later depressive symptoms.

The pattern of rsFC associations observed indicates that pubertal timing is linked to selective differences in connectivity across specific functional systems rather than to a uniform shift in connectivity across the connectome. Importantly, this pattern differed by sex. Network-pair variability in pubertal timing effects was substantially greater in females, indicating stronger heterogeneity across networks, whereas in males was smaller, suggesting more limited network-specific differences in connectivity related to pubertal timing. This pattern is consistent with the earlier pubertal onset and progression in females relative to males (53–55). Within the ABCD BL age range (9–11 years), many girls have progressed intro more advanced pubertal stages, whereas many boys remain in earlier stages of development (Fig. S4). Consequently, pubertal timing may capture more meaningful differences in biological maturation in females, while in males it may reflect more limited variability in underlying neuroendocrine processes.

Consistent with this interpretation, pubertal timing–rsFC associations were more pronounced and reproducible in females, particularly in longitudinal analyses. Although some cross-sectional network-pair associations were observed in males, these effects did not persist longitudinally, with no stable pubertal timing–rsFC_Y2_ associations detected. This suggests that pubertal timing assessed at ages 9–11 may not yet capture the developmental stage at which puberty-related differences in brain connectivity become detectable in boys. This aligns with developmental evidence indicating that neural and behavioral consequences of pubertal maturation often emerge later in males (53–55), potentially shifting detectable brain–puberty associations to slightly older ages.

In females, earlier pubertal timing was associated with a heterogeneous pattern of increases and decreases in rsFC across multiple large-scale functional systems. Negative associations were strongest for connections involving the BG, sensory–motor networks (e.g., AUD, SML), and association networks (e.g., VAN, SAL), suggesting reduced coupling between subcortical systems and distributed cortical networks involved in sensory processing and attention in earlier-maturing females relative to their same-age, same-sex peers. In contrast, positive associations involved interactions among sensory–motor systems (e.g., SCAN, SML, SMD) and association networks (e.g., VAN, FPN, DAN), suggesting stronger coupling between sensorimotor and attention systems. These findings are broadly consistent with prior work linking earlier pubertal maturation to heterogeneous alterations in large-scale functional connectivity, particularly involving subcortical–cortical interactions. For example, Vijayakumar et al. reported that pubertal timing was associated with reduced connectivity between limbic and striatal regions (i.e., amygdala and hippocampus) and multiple cortical networks, including salience-related, sensorimotor, ventral attention, and visual systems (39). More broadly, developmental studies suggest that pubertal maturation contributes to reorganization of cortico-subcortical connectivity beyond age-related effects (56). Consistent with this, prior work has shown both decreases and increases in connectivity between striatal or limbic regions and medial prefrontal or cingulate systems (57), with sex-differentiated trajectories such that advancing puberty in girls is associated with weaker DMN and DMN–control network connectivity, whereas boys show the opposite pattern (58). Together, these findings indicate that earlier pubertal timing in females is associated with a heterogeneous reorganization of large-scale functional brain networks, characterized by both increased and decreased connectivity across subcortical, sensory–motor, and association systems. This pattern suggests that earlier maturation may accelerate or alter the integration and differentiation of these networks, potentially reflecting earlier reorganization of connectivity between subcortical regulatory systems and cortical networks involved in attention and sensory processing.

In males, pubertal timing–rsFC associations were more limited, less heterogeneous, and weaker in magnitude across network pairs than those observed in females. Negative associations primarily involved the CEREB and interactions with sensory–motor and visual systems, whereas positive associations involved coupling among sensory–motor networks and higher-order association systems (e.g., CAN). This more limited pattern is consistent with prior developmental work suggesting that puberty-related changes in brain connectivity in boys are more circumscribed and centered on cerebellar and sensorimotor systems. Both longitudinal and cross-sectional rsFC studies have reported puberty-related changes in cerebellar–striatal and sensorimotor connectivity, including sex-moderated increases in putamen integration and alterations in cerebellar–sensorimotor hubs (11,56,59). Together, these findings suggest that, in males, the influence of pubertal timing on brain connectivity at this age is relatively constrained, primarily affecting motor and sensory systems. This pattern may reflect the later progression of pubertal maturation in boys, such that large-scale integration across distributed cortical networks emerges at later stages of development.

In longitudinal analyses, earlier pubertal timing predicted lower rsFC_Y2_ in females after adjusting for rsFC_BL_, indicating that pubertal timing was associated with subsequent changes in network organization during early adolescence. Compared with the cross-sectional findings, these longitudinal associations involved a smaller subset of network pairs but showed consistent effects across cross-validation folds. Notably, most longitudinal associations were negative, suggesting a general pattern of reduced connectivity over time among networks influenced by earlier pubertal maturation. These effects were concentrated in subcortical structures, particularly the BG and THAL, as well as sensory–motor systems and the VAN. Earlier pubertal maturation may therefore reflect earlier specialization or differentiation of large-scale functional brain networks, resulting in reduced connectivity among these systems during this developmental period. In contrast, no longitudinal associations between pubertal timing and rsFC_Y2_ were detected in males. This absence of effects may reflect the later onset and progression of pubertal maturation in boys, such that pubertal timing assessed at ages 9–11 may capture less meaningful variation in underlying biological maturation and therefore be less predictive of subsequent changes in brain connectivity.

Consistent with prior research (6,24,36,60–62), earlier pubertal timing was associated with higher depressive symptoms both concurrently and longitudinally. Youth who were more advanced in pubertal development reported higher depression_BL_, and pubertal timing predicted increases in depressive symptoms over time after accounting for prior symptom levels. These findings align with a substantial body of work demonstrating that earlier pubertal maturation is a robust risk factor for internalizing difficulties during adolescence (6,24,36,60–62). Although some studies have reported stronger associations in girls (6,29,63,64), the relationship between pubertal timing and depressive symptoms in the present sample did not significantly differ by sex. Together, these results reinforce the importance of pubertal timing as a developmental factor associated with emerging depressive symptoms during early adolescence.

Despite the associations between pubertal timing, rsFC, and depressive symptoms, rsFC_Y2_ did not mediate the relationship between pubertal timing and later depressive symptoms (depression_Y3_, controlling for depression_Y2_). These findings differ from those of Vijayakumar et al., who reported corticolimbic connectivity as a significant mediator linking pubertal timing and depressive symptoms in the ABCD Study (39). Several methodological differences may account for this discrepancy. In their study, pubertal timing, rsFC, and depressive symptoms were modeled concurrently by combining observations from both BL and Y2 assessments within the same analysis. Although this approach increases statistical power, it does not clearly separate predictor, mediator, and outcome in time and therefore mixes within- and between-person variation, limiting the ability to infer temporally ordered mediation effects. In contrast, the present longitudinal framework explicitly orders variables in time and controls for prior levels of rsFC, providing a more stringent test of whether pubertal timing predicts subsequent changes in brain connectivity that in turn relate to later depressive symptoms. Additionally, the broader age range (i.e., 9–11 and 10.5–13.5) in their study extends into slightly later developmental stages, when pubertal and neural changes may be more advanced. Finally, their analysis focused on corticolimbic connections, whereas the present study examined rsFC across the full connectome. While this broader approach enables a more comprehensive assessment of network-level associations, it may also be less sensitive to effects that are specific to corticolimbic circuits, potentially contributing to differences in observed mediation effects. These differences suggest that previously reported mediation effects may partly reflect concurrent associations or stable individual differences rather than longitudinal pathways linking pubertal timing, brain connectivity, and depressive symptoms during early adolescence.

The present study is one of the few investigations to examine associations among pubertal timing, whole-connectome rsFC, and depression during early adolescence in a large longitudinal sample. Nevertheless, some limitations should be considered. First, pubertal timing was derived from parent-reported measures rather than hormonal assessments, which may introduce measurement imprecision in estimating biological maturation. Future studies incorporating hormonal measures or multimodal assessments of pubertal development may help refine the characterization of puberty-related neural changes. Second, the age range examined here (9–11 years at BL) represents an early stage of pubertal development, particularly for males, potentially limiting detection of puberty-related neural changes emerging later in adolescence. Extending longitudinal analyses into later adolescent stages will therefore be important for determining whether puberty-related differences in brain connectivity become more pronounced as maturation progresses. Third, rsFC was examined at the network-pair level, which may obscure more localized or region-specific effects of pubertal development. Future studies combining network-level and region-level analyses may provide a more detailed understanding of how puberty influences functional brain organization. Finally, we did not directly examine potential contextual or psychological moderators such as family conflict, peer difficulties, or socioeconomic adversity, or co-occurring internalizing symptoms such as anxiety, which may shape how pubertal maturation relates to brain development and mental health risk. Future work incorporating such moderators could help identify the conditions under which earlier pubertal timing confers greater vulnerability to depression.

In conclusion, pubertal timing is associated with heterogeneous and sex-differentiated patterns of large-scale rsFC during early adolescence. Rather than reflecting a global shift in brain connectivity, earlier pubertal maturation was linked to selective differences across specific network systems, particularly involving subcortical and sensorimotor circuits in females. Although earlier pubertal timing was also associated with higher depressive symptoms, the alterations in rsFC alone did not appear to explain this relationship. These findings highlight pubertal timing as a key developmental factor shaping early brain functional network organization and underscore the need for future longitudinal work to clarify how puberty-related neural changes interact with environmental, and psychological factors to influence adolescent mental health.

## Acknowledgements

This work was supported by NIH grants NS140256 (EMG, NUFD), MH122066 (EMG, NUFD), MH121276 (EMG, NUFD), MH124567 (EMG, NUFD), NS129521 (EMG, NUFD), NS088590 (NUFD), MH134966 (TOL and EMG), MH129616 (TOL), MH121518 (SM), and U01DA041120-01 (D.M.B.); by the McDonnell Center for Systems Neuroscience (AM); by Michael J. Fox Foundation grant MJFF-026625 (EMG, NUFD); by the National Spasmodic Dysphonia Association (EMG); by the Taylor Family Foundation (TOL); by the Intellectual and Developmental Disabilities Research Center (NUFD); by the Kiwanis Foundation (NUFD); by the Washington University Hope Center for Neurological Disorders (EMG, BPK, NUFD); and by Mallinckrodt Institute of Radiology pilot funding (EMG, NUFD).

## Author Contributions

**Athanasia Metoki:** Conceptualization, Methodology, Validation, Formal analysis, Investigation, Data Curation, Writing – Original Draft, Writing – Review & Editing, Visualization, Supervision **Benjamin P. Kay:** Methodology, Writing – Review & Editing, **Roselyne Chauvin:** Writing – Review & Editing, **Evan M. Gordon:** Writing – Review & Editing, **Timothy O. Laumann:** Writing – Review & Editing, **Samuel R. Krimmel:** Writing – Review & Editing, **Scott Marek:** Writing – Review & Editing, **Anxu Wang:** Writing – Review & Editing, **Philip N. Cho:** Writing – Review & Editing, **Noah J. Baden:** Writing – Review & Editing, **Kristen M. Scheidter:** Writing – Review & Editing, **Julia Monk:** Writing – Review & Editing, **Nico U.F. Dosenbach:** Writing – Review & Editing, **Deanna M. Barch:** Methodology, Writing – Review & Editing, Supervision.

## References

1. Blakemore SJ, Burnett S, Dahl RE. The role of puberty in the developing adolescent brain. Hum Brain Mapp. 2010 Jun 1;31(6):926–33. doi:10.1002/HBM.21052 PubMed PMID: 20496383.

2. Herting MM, Sowell ER. Puberty and structural brain development in humans. Frontiers in Neuroendocrinology. Academic Press Inc.; 2017. p. 122–37. doi:10.1016/j.yfrne.2016.12.003 PubMed PMID: 28007528.

3. Juraska JM, Willing J. Pubertal onset as a critical transition for neural development and cognition. Brain Res. 2017;1654(Part B):87–94. doi:10.1016/j.brainres.2016.04.012

4. Pfeifer JH, Allen NB. Puberty Initiates Cascading Relationships Between Neurodevelopmental, Social, and Internalizing Processes Across Adolescence. Biol Psychiatry. 2021;89(2):99–108. doi:10.1016/j.biopsych.2020.09.002

5. Sanders AFP, Baum GL, Harms MP, Kandala S, Bookheimer SY, Dapretto M, et al. Developmental trajectories of cortical thickness by functional brain network: The roles of pubertal timing and socioeconomic status. Dev Cogn Neurosci. 2022;57:101145. doi:10.1016/j.dcn.2022.101145

6. MacSweeney N, Allardyce J, Edmondson-Stait A, Shen X, Casey H, Chan SWY, et al. The role of brain structure in the association between pubertal timing and depression risk in an early adolescent sample (the ABCD Study®): A registered report. Dev Cogn Neurosci. 2023 Apr 1;60:101223. doi:10.1016/J.DCN.2023.101223 PubMed PMID: 36870214.

7. Cai L, Dong Q, Niu H. The development of functional network organization in early childhood and early adolescence: A resting-state fNIRS study [Internet]. 2018. doi:10.1016/j.dcn.2018.03.003

8. İçer S. Functional connectivity differences in brain networks from childhood to youth. Int J Imaging Syst Technol. 2020 Mar 1;30(1):75–91. doi:10.1002/IMA.22366

9. Tapia Medina MG, Cosío-Guirado R, Peró-Cebollero M, Cañete-Massé C, Villuendas-González ER, Guàrdia-Olmos J. The clinical relevance of healthy neurodevelopmental connectivity in childhood and adolescence: a meta-analysis of resting-state fMRI. Front Neurosci. 2025 Jun 26;19:1576932. doi:10.3389/FNINS.2025.1576932/TEXT

10. Teeuw J, Brouwer RM, Ao J, Guimar∼ Aes C, Poft, Brandner P, Koenis MG, et al. Genetic and environmental influences on functional connectivity within and between canonical cortical resting-state networks throughout adolescent development in boys and girls [Internet]. 2019. doi:10.1016/j.neuroimage.2019.116073

11. Sanders AFP, Harms MP, Kandala S, Marek S, Somerville LH, Bookheimer SY, et al. Age-related differences in resting-state functional connectivity from childhood to adolescence. Cerebral Cortex. 2023;33:6928–42. doi:10.1093/cercor/bhad011

12. Kwong ASF, Manley D, Timpson NJ, Pearson RM, Heron J, Sallis H, et al. Identifying Critical Points of Trajectories of Depressive Symptoms from Childhood to Young Adulthood. J Youth Adolesc. 2019;48:815–27. doi:10.1007/s10964-018-0976-5

13. Lu B, Lin L, Su X. Global burden of depression or depressive symptoms in children and adolescents: A systematic review and meta-analysis. J Affect Disord. 2024;354:553–62. doi:10.1016/j.jad.2024.03.074

14. Shorey S, Ng ED, Wong CHJ. Global prevalence of depression and elevated depressive symptoms among adolescents: A systematic review and meta-analysis. British Journal of Clinical Psychology. 2022 Jun 1;61(2):287–305. doi:10.1111/BJC.12333 PubMed PMID: 34569066.

15. Thapar A, Collishaw S, Pine DS, Thapar AK. Depression in adolescence. The Lancet. 2012 Mar;379(9820):1056–67. doi:10.1016/S0140-6736(11)60871-4

16. Salk RH, Hyde JS, Abramson LY. Gender Differences in Depression in Representative National Samples: Meta-Analyses of Diagnoses and Symptoms. Psychol Bull. 2017 Aug;143(8):783–822. doi:10.1037/bul0000102

17. Hankin BL, Young JF, Abela JRZ, Smolen A, Jenness JL, Gulley LD, et al. Depression From Childhood Into Late Adolescence: Influence of Gender, Development, Genetic Susceptibility, and Peer Stress. J Abnorm Psychol. 2015;124(4):803–16. doi:10.1037/abn0000089

18. Richburg AG, Kelly DP, Davis-Kean PPE, Richburg A. Depression, Anxiety, and Pubertal Timing: Current Research and Future Directions. University of Michigan Undergraduate Research Journal. 2021;15. doi:10.3998/UMURJ.1383

19. Kaltiala-Heino R, Kosunen E, Rimpelä M. Pubertal timing, sexual behaviour and self-reported depression in middle adolescence. J Adolesc. 2003;26(5):531–45. doi:10.1016/S0140-1971(03)00053-8 PubMed PMID: 12972267.

20. Hamlat EJ, Snyder HR, Young JF, Hankin BL. Pubertal Timing as a Transdiagnostic Risk for Psychopathology in Youth. Clinical Psychological Science. 2019 May 1;7(3):411–29. doi:10.1177/2167702618810518/FORMAT/EPUB

21. Graber JA. Pubertal timing and the development of psychopathology in adolescence and beyond. Hormones and Behavior. 2013. p. 262–9. doi:10.1016/j.yhbeh.2013.04.003 PubMed PMID: 23998670.

22. Angold A, Costello EJ, Worthman CM. Puberty and depression: the roles of age, pubertal status and pubertal timing. Psychol Med. 1998;28:61. doi:10.1017/S003329179700593X

23. Ge X, Natsuaki MN. In search of explanations for early pubertal timing effects on developmental psychopathology. Curr Dir Psychol Sci. 2009;18(6):327–31. doi:10.1111/J.1467-8721.2009.01661.X/FORMAT/EPUB

24. Hamlat EJ, McCormick KC, Young JF, Hankin BL. Early pubertal timing predicts onset and recurrence of depressive episodes in boys and girls. Journal of Child Psychology and Psychiatry. 2020 Nov 1;61(11):1266–74. doi:10.1111/JCPP.13198 PubMed PMID: 32017111.

25. Benoit A, Lacourse E, Claes M. Pubertal timing and depressive symptoms in late adolescence: The moderating role of individual, peer, and parental factors. Dev Psychopathol. 2013;25:455–71. doi:10.1017/S0954579412001174

26. Rudolph KD, Troop-Gordon W, Lambert SF, Natsuaki MN. Long-term consequences of pubertal timing for youth depression: Identifying personal and contextual pathways of risk. Development and Psyshopathology. 2014;26:1423–44. doi:10.1017/S0954579414001126

27. Vijayakumar N, Whittle S. A systematic review into the role of pubertal timing and the social environment in adolescent mental health problems. Clin Psychol Rev. 2023;102:102282. doi:10.1016/j.cpr.2023.102282

28. Barendse MEA, Byrne ML, Flournoy JC, McNeilly EA, Guazzelli Williamson V, Barrett AMY, et al. Multimethod assessment of pubertal timing and associations with internalizing psychopathology in early adolescent girls. Journal of Psychopathology and Clinical Science. 2022 Jan;131(1):14–25. doi:10.1037/ABN0000721

29. Galvao TF, Silva MT, Zimmermann IR, Souza KM, Martins SS, Pereira MG. Pubertal timing in girls and depression: A systematic review. Journal of Affective Disorders. 2014. p. 13–9. doi:10.1016/j.jad.2013.10.034 PubMed PMID: 24274962.

30. Mendle J, Harden KP, Brooks-Gunn J, Graber JA. Development’s tortoise and hare: Pubertal timing, pubertal tempo, and depressive symptoms in boys and girls. Dev Psychol. 2010 Sep;46(5):1341–53. doi:10.1037/a0020205 PubMed PMID: 20822243.

31. Marceau K, Ram N, Houts RM, Grimm KJ, Susman EJ. Individual Differences in Boys’ and Girls’ Timing and Tempo of Puberty: Modeling Development With Nonlinear Growth Models. Dev Psychol. 2011 Sep;47(5):1389–409. doi:10.1037/a0023838 PubMed PMID: 21639623.

32. Ullsperger JM, Nikolas MA. A Meta-Analytic Review of the Association Between Pubertal Timing and Psychopathology in Adolescence: Are There Sex Differences in Risk? Psychol Bull. 2017;143(9):903–38. doi:10.1037/bul0000106.supp

33. Dinkelbach L, Peters T, Grasemann C, Hinney A, Hirtz R. The causal role of male pubertal timing for the development of externalizing and internalizing traits: results from Mendelian randomization studies. medRxiv. 2024 May 31;2024.05.30.24308257. doi:10.1101/2024.05.30.24308257

34. Hamilton JL, Hamlat EJ, Stange JP, Abramson LY, Alloy LB. Pubertal timing and vulnerabilities to depression in early adolescence: Differential pathways to depressive symptoms by sex. J Adolesc. 2014 Feb;37(2):165–74. doi:10.1016/j.adolescence.2013.11.010 PubMed PMID: 24439622.

35. Hamlat EJ, Stange JP, Abramson LY, Alloy LB. Early pubertal timing as a vulnerability to depression symptoms: Differential effects of race and sex. J Abnorm Child Psychol. 2014 Sep 8;42(4):527–38. doi:10.1007/S10802-013-9798-9/FIGURES/2 PubMed PMID: 24014162.

36. Copeland WE, Worthman C, Shanahan L, Costello EJ, Angold A. Early Pubertal Timing and Testosterone associated with higher levels of Adolescent Depression in Females. J Am Acad Child Adolesc Psychiatry. 2019 Dec 1;58(12):1197. doi:10.1016/J.JAAC.2019.02.007 PubMed PMID: 30768421.

37. Mendle J, Ferrero J. Detrimental psychological outcomes associated with pubertal timing in adolescent boys. Developmental Review. 2012. p. 49–66. doi:10.1016/j.dr.2011.11.001

38. Negriff S, Susman EJ. Pubertal Timing, Depression, and Externalizing Problems: A Framework, Review, and Examination of Gender Differences. Journal of Research on Adolescence. 2011 Sep 1;21(3):717–46. doi:10.1111/J.1532-7795.2010.00708.X

39. Vijayakumar N, Whittle S, Silk TJ. Corticolimbic connectivity mediates the relationship between pubertal timing and mental health problems. Psychol Med. 2023 Dec 2;53(16):7655–65. doi:10.1017/S0033291723001472

40. Garavan H, Bartsch H, Conway K, Decastro A, Goldstein RZ, Heeringa S, et al. Recruiting the ABCD sample: Design considerations and procedures. Dev Cogn Neurosci. 2018 Aug 1;32:16–22. doi:10.1016/J.DCN.2018.04.004 PubMed PMID: 29703560.

41. Colich NL, Hanford LC, Weissman DG, Allen NB, Shirtcliff EA, Lengua LJ, et al. Childhood trauma, earlier pubertal timing, and psychopathology in adolescence: The role of corticolimbic development. Dev Cogn Neurosci. 2023 Feb 1;59:101187. doi:10.1016/J.DCN.2022.101187 PubMed PMID: 36640624.

42. Chahal R, Marek S, Vilgis V, Weissman D, Hastings P, Robins R, et al. Neural connectivity mechanisms linking off-time pubertal development and depression risk in adolescence. J Clin Transl Sci. 2019 Mar;3(1):17–8. doi:10.1017/cts.2019.42

43. Marek S, Tervo-Clemmens B, Calabro FJ, Montez DF, Kay BP, Hatoum AS, et al. Reproducible brain-wide association studies require thousands of individuals. Nature. 2022 Mar 16;603(7902):654–60. doi:10.1038/s41586-022-04492-9 PubMed PMID: 35296861.

44. Casey BJ, Cannonier T, Conley MI, Cohen AO, Barch DM, Heitzeg MM, et al. The Adolescent Brain Cognitive Development (ABCD) study: Imaging acquisition across 21 sites. Developmental Cognitive Neuroscience. Elsevier Ltd; 2018. p. 43–54. doi:10.1016/j.dcn.2018.03.001 PubMed PMID: 29567376.

45. Gordon EM, Laumann TO, Adeyemo B, Huckins JF, Kelley WM, Petersen SE. Generation and Evaluation of a Cortical Area Parcellation from Resting-State Correlations. Cerebral Cortex. 2016 Jan 1;26(1):288–303. doi:10.1093/CERCOR/BHU239 PubMed PMID: 25316338.

46. Seitzman BA, Gratton C, Marek S, Raut R V., Dosenbach NUF, Schlaggar BL, et al. A set of functionally-defined brain regions with improved representation of the subcortex and cerebellum. Neuroimage. 2020 Feb 1;206:116290. doi:10.1016/J.NEUROIMAGE.2019.116290 PubMed PMID: 31634545.

47. Fisher RA. Frequency Distribution of the Values of the Correlation Coefficient in Samples from an Indefinitely Large Population. Biometrika. 1915 May;10(4):507. doi:10.2307/2331838

48. Dehestani N, Vijayakumar N, Ball G, Mansour L S, Whittle S, Silk TJ. “Puberty age gap”: new method of assessing pubertal timing and its association with mental health problems. Mol Psychiatry. 2023 Dec 5;1–8. doi:10.1038/s41380-023-02316-4

49. Petersen AC, Crockett L, Richards M, Boxer A. A self-report measure of pubertal status: Reliability, validity, and initial norms. J Youth Adolesc. 1988 Apr;17(2):117–33. doi:10.1007/BF01537962 PubMed PMID: 24277579.

50. Schlossberger NM, Turner RA, Irwin CE. Validity of self-report of pubertal maturation in early adolescents. Journal of Adolescent Health. 1992 Mar 1;13(2):109–13. doi:10.1016/1054-139X(92)90075-M PubMed PMID: 1627576.

51. Achenbach TM, Rescorla LA. Manual for the ASEBA preschool forms and profiles. Burlington, VT: University of Vermont, Research center for children, youth, & families. 2000.

52. Thurber S, Sheehan WP. Note on truncated T scores in discrepancy studies with the Child Behavior Checklist and Youth Self Report. Archives of Assessment Psychology [Internet]. 2012 Oct 17 [cited 2024 Oct 11];2(1):73–80. Available from: http://www.assessmentpsychologyboard.org/journal/index.php/AAP/article/view/39

53. Farello G, Altieri C, Cutini M, Pozzobon G, Verrotti A. Review of the literature on current changes in the timing of pubertal development and the incomplete forms of early puberty. Front Pediatr. 2019;7(MAR). doi:10.3389/FPED.2019.00147/FULL

54. Omary A, Curtis | Mark, Cheng TW, Mair P, Shirtcliff EA, Barch DM, et al. Multimodal Measurement of Pubertal Development: Stage, Timing, Tempo, and Hormones. Child Dev. 2025;0:1–20. doi:10.1111/cdev.14220

55. Beck D, Ferschmann L, MacSweeney N, Norbom LB, Wiker T, Aksnes E, et al. Puberty differentially predicts brain maturation in male and female youth: A longitudinal ABCD Study. Dev Cogn Neurosci. 2023 Jun 1;61. doi:10.1016/j.dcn.2023.101261 PubMed PMID: 37295068.

56. van Duijvenvoorde ACK, Westhoff B, de Vos F, Wierenga LM, Crone EA. A three-wave longitudinal study of subcortical–cortical resting-state connectivity in adolescence: Testing age- and puberty-related changes. Hum Brain Mapp. 2019;40(13):3769–83. doi:10.1002/hbm.24630 PubMed PMID: 31099959.

57. Ojha A, Parr AC, Foran W, Calabro FJ, Luna B. Puberty contributes to adolescent development of fronto-striatal functional connectivity supporting inhibitory control. Dev Cogn Neurosci. 2022;58:101183. doi:10.1016/j.dcn.2022.101183

58. Ernst M, Benson B, Artiges E, Gorka AX, Lemaitre H, Lago T, et al. Pubertal maturation and sex effects on the default-mode network connectivity implicated in mood dysregulation. Transl Psychiatry. 2019;103(9). doi:10.1038/s41398-019-0433-6

59. Pacheco-Blas L, González-González G, Ortega-Aguilar A. Sex-Related Variations in the Brain Motor-Network Connectivity at Rest during Puberty. Applied Sciences. 2023;23(10006):1–22. doi:10.3390/app131810006

60. Dahl Askelund A, Wootton RE, Torvik FA, Lawn RB, Ask H, Corfield EC, et al. Assessing causal links between age at menarche and adolescent mental health: a Mendelian randomisation study. BMC Med. 2024;21(155):1–27. doi:10.1186/s12916-024-03361-8

61. McGuire TC, McCormick KC, Koch MK, Mendle J. Pubertal maturation and trajectories of depression during early adolescence. Front Psychol. 2019 Jun 12;10(JUN):430978. doi:10.3389/FPSYG.2019.01362/BIBTEX

62. Prince C, Joinson C, Kwong ASF, Fraser A, Heron J. The relationship between timing of onset of menarche and depressive symptoms from adolescence to adulthood. Epidemiol Psychiatr Sci. 2023;32:e60. doi:10.1017/S2045796023000707

63. Alloy LB, Hamilton JL, Hamlat EJ, Abramson LY. Pubertal development, emotion regulatory styles, and the emergence of sex differences in internalizing disorders and symptoms in adolescence. Clinical Psychological Science. 2016 Sep 1;4(5):867–81. doi:10.1177/2167702616643008/SUPPL_FILE/SUPPL-MATERIAL.PDF PubMed PMID: 27747141.

64. Conley CS, Rudolph KD. The emerging sex difference in adolescent depression: Interacting contributions of puberty and peer stress. Dev Psychopathol. 2009;21(2):593–620. doi:10.1017/S0954579409000327 PubMed PMID: 19338700.

