## Supplemental_Information for "Functional brain network correlates of pubertal timing and depressive symptoms in preadolescence"

Metoki et al.

### Methods and Materials

#### Participants

Data for the present study were drawn from the Adolescent Brain Cognitive Development (ABCD) Study curated annual release 5.1, which includes data from the BL, Y2, and Y3 assessments (https://data-archive.nimh.nih.gov/abcd). The ABCD Study is an ongoing 10-year longitudinal study of 11,875 children enrolled at ages 9-10 years from 21 sites across the United States. The analytic sample for the present study included multiple nested samples depending on data availability across behavioral and neuroimaging modalities. A total of 10,957 children (5,243 females) had completed behavioral assessments at BL and were included in the derivation of the pubertal timing measure. Of these, 9,220 children (4,384 females) had complete behavioral data across all three assessment waves (BL, Y2, and Y3). The primary neuroimaging analytic sample comprised 7,594 children (3,682 females) with usable resting-state fMRI data at BL. For the longitudinal mediation analyses, the sample was further restricted to participants with available imaging and behavioral data at all three waves, yielding a final sample of 3,136 youth (1,465 females) (Fig. 1A; Fig. S1; Table S1). Informed written consent was obtained from parents or legal guardians, and written assent was obtained from the children. All study procedures were approved by the Institutional Review Board at the University of California, San Diego, and by the IRBs of each participating site.

#### fMRI Data Acquisition and Processing

Resting-state fMRI data were obtained from the ABCD Study’s BL and Y2 assessments. Twenty minutes (4 × 5 min runs) of eyes-open (passive crosshair viewing) resting-state fMRI data were collected to ensure at least 5 min of low-motion data. All resting-state fMRI scans were acquired using a gradient-echo EPI sequence (TR = 800 ms, TE = 30 ms, flip angle = 90◦, voxel size = 2.4 mm3, 60 slices). Head motion was monitored online using the Framewise Integrated Real-time MRI Monitor (FIRMM) software at Siemens sites (1).

The image processing stream has been detailed elsewhere (2,3). Briefly, the ABCD pipeline comprises six stages: (1) PreFreeSurfer, which normalizes anatomical data; (2) FreeSurfer, which constructs cortical surfaces from the normalized anatomical data; (3) PostFreeSurfer, which converts outputs from FreeSurfer to CIFTIs and transforms the volumes to a standard volume space using ANTs nonlinear registration; (4) “Vol” stage, which performs the atlas transformation, mean field distortion correction, and resampling to 2 mm isotropic voxels in a single step using FSL’s applywarp tool; (5) “Surf” stage, which projects the volumetric functional data onto the surface; and (6) “DCANBOLDproc”, which performs functional connectivity processing.

DCANBOLDproc includes a respiratory filter to improve FD estimates calculated in the “vol” stage. Temporal masks were created to flag motion-contaminated frames using the improved FD estimates (4). Frames with FD > 0.20 mm were flagged as motion contaminated and removed. After computing the temporal masks for high motion frame censoring, the data were processed with the following steps: (i) demeaning and detrending, (ii) interpolation across censored frames using least squares spectral estimation of the values at censored frames (5) so that continuous data can be (iii) denoised via a general linear model (GLM) including whole brain, ventricular, and white matter signals, as well as their derivatives. Denoised data were then passed through (iv) a band-pass filter (0.008 Hz < f < 0.10 Hz) without re-introducing nuisance signals (6) or contaminating frames near high motion frames (7).

A region-of-interest (ROI) approach was employed, incorporating a total of 394 parcels: 333 cortical parcels defined by Gordon et al. (8) and 61 subcortical parcels from Seitzman et al. (9). For each participant, resting-state functional connectivity (rsFC) timecourses were extracted from these ROIs. The 333 cortical and 61 subcortical ROIs were averaged according to their network or anatomical grouping to generate timecourses for higher-order regions. Specifically, the cortical ROIs were organized into 13 canonical large-scale networks: default mode (DMN), somato-cognitive action (SCAN), visual (VIS), frontoparietal (FPN), dorsal attention (DAN), ventral attention (VAN), salience (SAL), action mode (AMN), sensorimotor dorsal (SMD), somatomotor lateral (SML), auditory (AUD), parietal memory (PMN), and context association (CAN). The subcortical ROIs were grouped into the following five anatomical structures: hippocampus (HIPP), amygdala (AMYG), basal ganglia (BG), thalamus (THAL), and cerebellum (CEREB). Pearson correlation coefficients were computed between the timecourse of each network/structure and every other network/structure, resulting in a total of 171 unique connections per participant. These correlation values were then normalized using Fisher’s r-to-z transformation (10).

#### Pubertal timing

To estimate pubertal timing, we implemented a generalized additive model (GAM) with nested 10-fold cross-validation, using MATLAB’s (R2023b; version 23.2.0; Update 10) (11) “fitrgam” function. In each iteration, the model was trained on 90% of the data and tested on the remaining 10%, with an inner loop performing grid search-based hyperparameter tuning. Model training was conducted on BL data only. To reduce confounding from psychopathology, the model was trained on “typically developing” participants (Fig. 1A), defined by parent-reported Child Behavior Checklist (CBCL) scores below 60 across DSM-oriented and broadband scales (12). To further ensure the independence of training observations, only one child per family was included in this training subset. The final model was then applied to the full sample to derive individual-level pubertal timing estimates.

#### Statistical Analyses

##### Baseline associations between pubertal timing and rsFC

To characterize how pubertal timing relates to large-scale brain organization at BL, we examined associations between pubertal timing rsFC across network pairs using complementary frequentist and Bayesian multilevel approaches. Frequentist models evaluated global effects and variance components, whereas Bayesian multilevel models provided network-pair–specific posterior estimates with principled uncertainty quantification and partial pooling. Together, these approaches allowed us to test overall associations and characterize connection-level heterogeneity. Pubertal timing and age were standardized prior to analysis, and singleton families were collapsed to stabilize estimation of family-level random effects.

Given strong biological motivation for sex differences in pubertal development and brain maturation (13–16), we evaluated whether associations between pubertal timing and rsFC were adequately characterized by a shared analytic framework across sexes or required sex-specific modeling. We fit a pooled linear mixed-effects model, using “lme4” (v1.1.37) (17) in R (v4.5.0) (18), including a pubertal timing × sex interaction and network-pair–specific random slopes for pubertal timing, with random intercepts for site, family, and participant. Sex-specific heterogeneity was evaluated by comparing the pooled model to an extended pooled model which allowed female-specific deviations using likelihood ratio tests, the Akaike Information Criterion (AIC; (19)) and Bayesian Information Criterion (BIC; (20)). Lower information criterion values indicated improved model fit. To aid interpretation, sex-stratified linear mixed-effects models were subsequently fit using identical model specifications. For each sex, we began with a maximal random-effects structure, evaluated variance components, and subsequently adopted reduced models that retained supported random effects (Equation S1). These sex-specific models were used to estimate population-level associations between pubertal timing and rsFC_BL_ within each sex.

Equation S1. Cross-sectional pubertal timing–rsFC mixed-effects model

$$\mathrm{rsFC}_{\mathrm{BL}} \sim Pubertal timing + \mathrm{Age}_{\mathrm{BL}}+ (0 + Pubertal timing || Network pair ID) + (1 | Site) + (1 | Family ID) + (1 | Participant)$$

Finally, to obtain network-pair–specific uncertainty estimates, Bayesian multilevel models were fit using “brms” (v2.23.0) (21–23) in R, specifying the same fixed- and random-effects structure as in the corresponding frequentist models. Posterior distributions of pubertal timing effects were extracted for each network pair, and credible intervals were used to summarize the magnitude, direction, and uncertainty of pubertal timing–rsFC_BL_ associations under a partially pooled hierarchical framework. Because network pairs were modeled as levels within the same hierarchical model, estimates were regularized via partial pooling (hierarchical shrinkage), and no additional multiple-comparison corrections were applied (24).

##### Longitudinal pubertal timing effects on rsFC

To examine whether pubertal timing was associated with longitudinal differences in rsFC, we conducted sex-stratified analyses. Within each sex, rsFC at the 2-year follow-up (rsFC_Y2_) was modeled as a function of pubertal timing while adjusting for age at Y2 (age_Y2_) and rsFC_BL_. We first fit a linear mixed-effects model with network-pair–specific random slopes for pubertal timing and random intercepts for site, family, and participant (Equation S2). Including rsFC_BL_ as a covariate allowed us to isolate pubertal-timing–related differences in rsFC_Y2_ above and beyond the effects of rsFC_BL_, while allowing associations to vary across rsFC network pairs. This analysis provided an initial estimate of the magnitude and variability of pubertal-timing effects across the connectome.

Equation S2. Longitudinal pubertal timing–rsFC mixed-effects model

$$\mathrm{rsFC}_{Y2} \sim Pubertal timing + \mathrm{Age}_{Y2}+ \mathrm{rsFC}_{Y1}+ (0 + Pubertal timing || Network pair ID) + (1 | Site) + (1 | Family ID) + (1 | Participant)$$

To identify rsFC network pairs showing stable longitudinal associations with pubertal timing for inclusion in subsequent mediation analyses, we implemented a 5-fold, family-level cross-validation procedure using Bayesian multilevel modeling. This approach ensured that only reproducible longitudinal effects were carried forward into mediation models, enhancing robustness. Families were assigned to a single fold to prevent related individuals from being split across training and test sets, with fold assignment implemented in R using “caret” (v7.0.1) (25). A random-search balancing algorithm minimized between-fold deviations from the overall sample on key variables, including pubertal timing and age_Y2_ (standardized mean differences), as well as race/ethnicity and site (L1 distances between categorical distributions). Within each fold, model fitting and network-pair selection were performed using training data only. Prior to modeling, pubertal timing, age_Y2_, and all rsFC measures were standardized within the training set, and the resulting scaling parameters were applied only to the corresponding training data. Bayesian multilevel models mirroring the longitudinal mixed-effects models (Equation S2) were then fit within each training set. Posterior distributions of pubertal timing effects were extracted for each network pair, and pairs with 95% credible intervals excluding zero were identified. Selection stability was defined as selection in ≥3 of 5 training folds with a consistent direction of effect. Network pairs meeting this stability criterion were carried forward for mediation analyses in the full sample. Final posterior estimates were obtained by refitting the model in the full sample.

##### Associations between pubertal timing and depressive symptoms, and network-specific mediation of depressive symptoms

To evaluate associations between pubertal timing and depressive symptoms, we conducted both cross-sectional and longitudinal mixed-effects analyses. First, we examined whether pubertal timing was associated with depression_BL_ by fitting linear mixed-effects models with pubertal timing, sex, and their interaction as predictors, adjusting for age at BL (age_BL_) (Equation S3). Random intercepts for site and family accounted for non-independence of observations.

Equation S3. Cross-sectional pubertal timing–depression mixed-effects model

$$\mathrm{depression}_{\mathrm{BL}}\sim Pubertal timing*Sex + \mathrm{Age}_{\mathrm{BL}}+ (1 | Site) + (1 | Family ID)$$

We then examined whether pubertal timing predicted later depressive symptoms by fitting longitudinal linear mixed-effects models. Depressive symptoms at Y3 (depression_Y3_) were modeled as a function of pubertal timing, sex (and their interaction), and depressive symptoms at Y2 (depression_Y2_) to account for prior symptom levels (Equation S4). Age at Y3 (Age_Y3_) was included as a covariate, along with random intercepts for site and family. Together, these analyses assessed both contemporaneous and prospective associations between pubertal timing and depressive symptoms.

Equation S4. Longitudinal pubertal timing–depression mixed-effects model

$$\mathrm{depression}_{Y3}\sim Pubertal timing*Sex + \mathrm{depression}_{Y2}+\mathrm{Age}_{Y3}+ (1 | Site) + (1 | Family ID)$$

Finally, we tested a longitudinal, network-specific mediation framework using Bayesian multilevel mediation analysis. The model included a mediator path (Path A), in which pubertal timing predicted rsFC_Y2_, controlling for rsFC_BL_ and age_Y2_, and an outcome path (Path B), in which rsFC_Y2_ predicted depression_Y3_, adjusting for depression_Y2_ and age_Y3_. Path A was modeled using a Gaussian distribution, whereas Path B a negative binomial distribution with a log link to account for overdispersion in CBCL Withdrawn/Depressed scores. Both paths included network-pair–specific random slopes, allowing mediation effects to vary across large-scale network pairs.

Weakly informative priors were specified for all fixed and random effects to regularize estimation while minimizing undue influence on posterior inference (26). Models were fit using four Markov chain Monte Carlo (MCMC) chains with 4,000 iterations per chain, including 2,000 warm-up iterations, yielding 8,000 post–warm-up samples per model. Sampling was conducted with a high target acceptance probability during Hamiltonian adaptation (adapt_delta = 0.99) and increased tree depth (max_treedepth = 15) to promote stable estimation. Model convergence was assessed using R̂ statistics (<1.05 for all parameters) and posterior predictive checks.

Posterior samples were used to compute indirect effects for each network pair as the product of the corresponding Path A and Path B coefficients. Direct effects reflected the association between pubertal timing and depressive symptoms after accounting for rsFC, and total effects were calculated as the sum of direct and indirect effects. All effects were summarized using posterior medians and equal-tailed 95% credible intervals.

Supplementary analyses included body mass index (BMI) and parental mood in the outcome model (Path B), given evidence that BMI is associated with early pubertal timing (27) and depression (28), and that parental mood may influence parental reports of child psychopathology (29). These variables were not included in the primary analyses to minimize sample size reduction due to missing data (*n* = 374).

##### Other statistical analyses

To characterize the sample and assess potential sex-related differences in key demographic and developmental variables that could influence subsequent analyses, we conducted descriptive comparisons between males and females. Independent-samples t-tests were used to compare age, which was approximately normally distributed, whereas Wilcoxon rank-sum tests were used to compare pubertal timing and CBCL Withdrawn/Depressed scores at BL, Y2, and Y3 due to non-normal and heteroskedastic distributions.

### Supplementary Figures

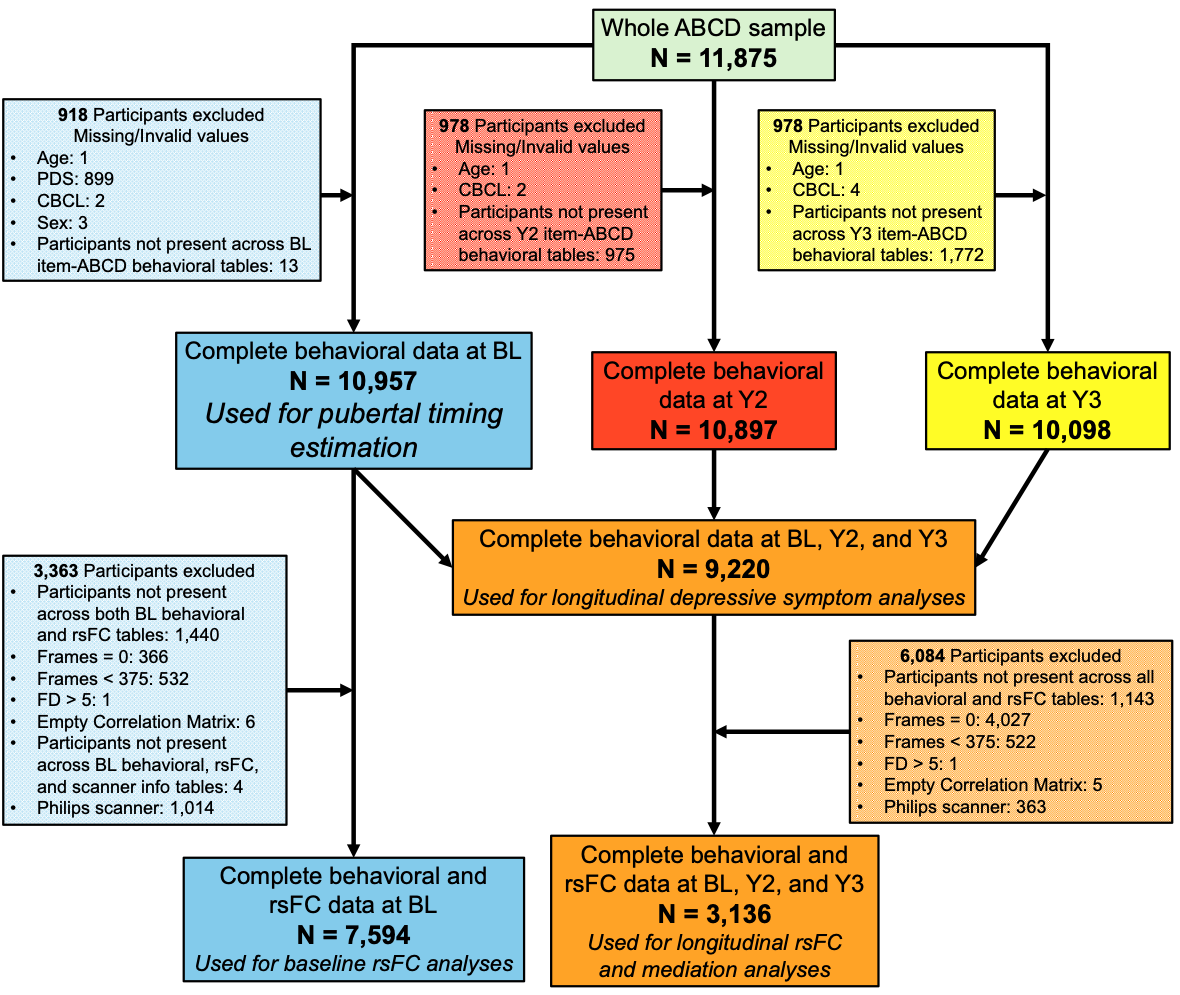

**Figure S1. Sample derivation.** Flow diagram illustrating sample derivation. The full baseline (BL) cohort (n = 11,875) was sequentially restricted to participants with complete behavioral data at BL, complete behavioral data across BL and follow-ups, and complete resting-state functional connectivity (rsFC) data at BL and longitudinally. Subsamples used for pubertal timing calculation, depressive symptom analyses, cross-sectional rsFC analyses, and longitudinal mediation models are indicated. BL: baseline; Y2: 2-year follow-up; Y3: 3-year follow-up assessment

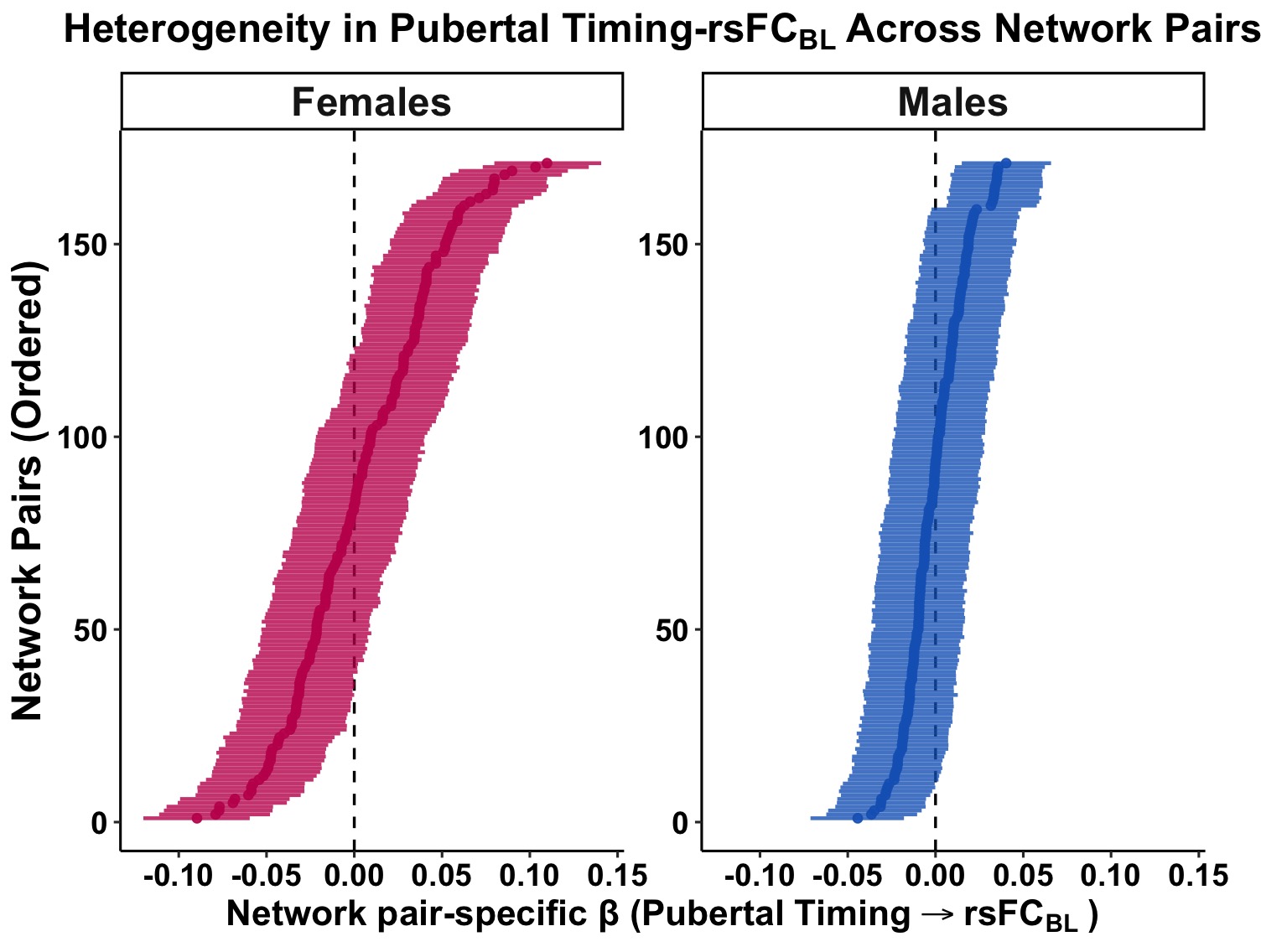

**Figure S2. Network-pair heterogeneity in associations between pubertal timing and baseline resting-state functional connectivity (rsFC) in females and males**. Points represent posterior median network pair-specific β estimates for associations between pubertal timing and rsFC at baseline, and horizontal lines indicate 95% credible intervals. Network pairs are ordered within each sex from the most negative to the most positive association. Negative β values indicate lower rsFC with earlier pubertal timing, whereas positive β values indicate higher rsFC with earlier pubertal timing. The dashed vertical line indicates β = 0. BL: baseline assessment; rsFC: resting-state functional connectivity.

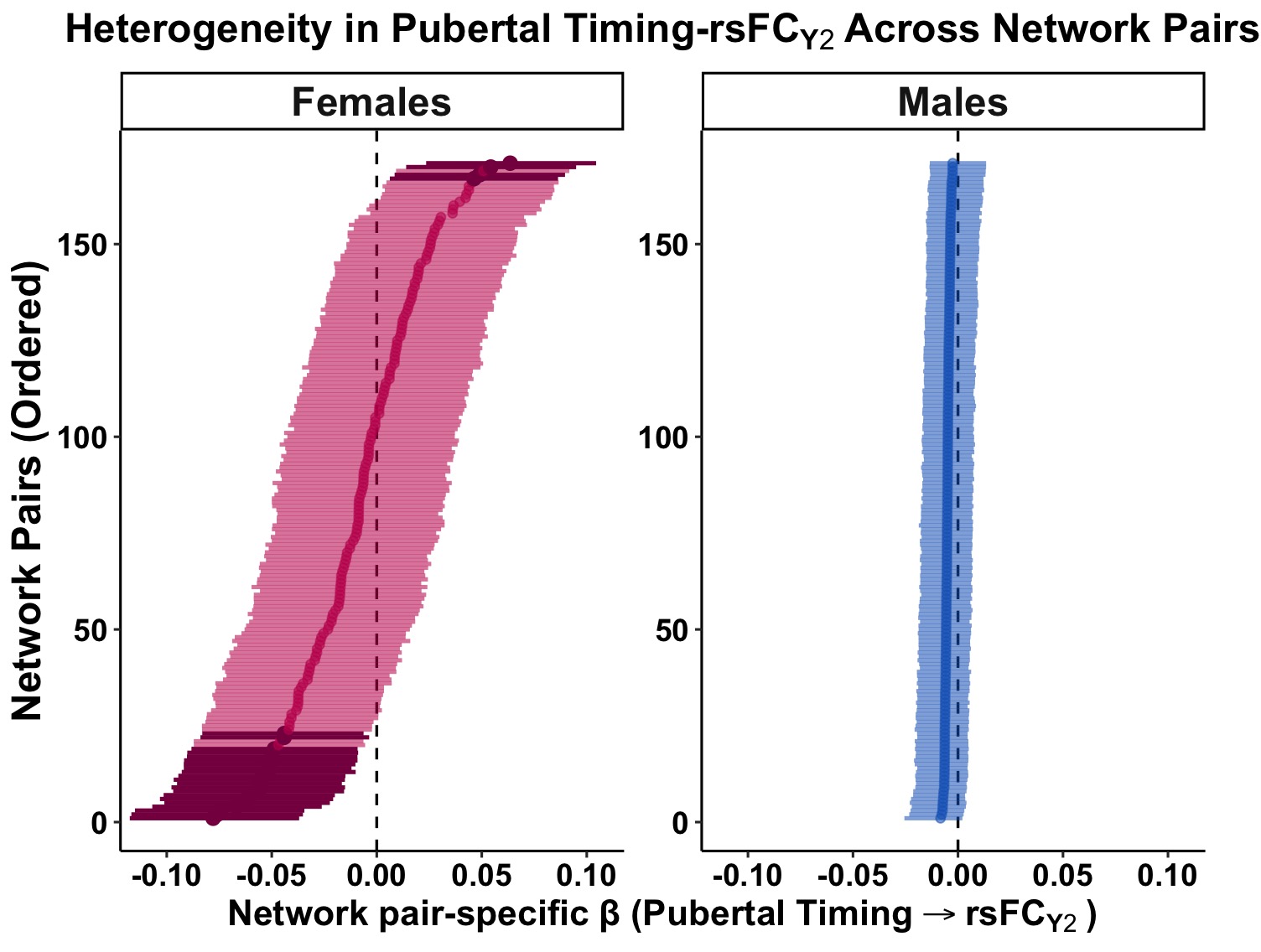

**Figure S3. Network-pair heterogeneity in associations between pubertal timing and 2-year follow-up resting-state functional connectivity**. Points represent posterior median network pair-specific β estimates for associations between pubertal timing and rsFC at baseline, and horizontal lines indicate 95% credible intervals. Darker points and intervals indicate network pairs showing stable associations across cross-validation folds (≥3/5 folds with consistent direction). Network pairs are ordered within each sex from the most negative to the most positive association. Negative β values indicate lower rsFC with earlier pubertal timing, whereas positive β values indicate higher rsFC with earlier pubertal timing. The dashed vertical line indicates β = 0. Y2: 2-year follow-up assessment; rsFC: resting-state functional connectivity.

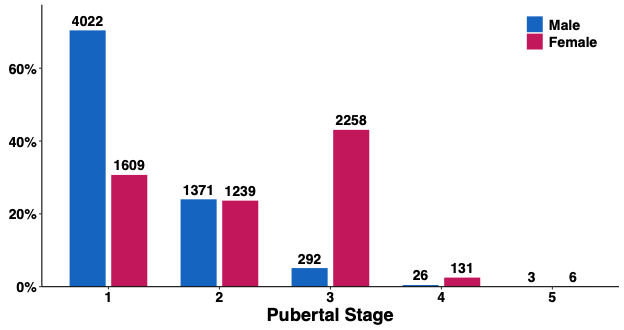

**Figure S4. Distribution of pubertal stage at baseline by sex**. Bars show the proportion of participants within each sex across pubertal stages at baseline (ages 9–11 years). Females are more likely to be in more advanced pubertal stages, whereas males are more likely to remain in earlier stages of development. Numbers above bars indicate the number of participants in each category.

### Supplementary Tables

**Table S1. Sample characteristics and sex differences in age, pubertal timing, and depressive symptoms.**

| **Variable** | **Males**  **(n = 4,836)** | **Females**  **(n = 4,384)** | **Test Statistic** | **p value** |
| --- | --- | --- | --- | --- |
| **Age (BL, months)**  Mean ± SD | 119 ± 7.5 | 119 ± 7.5 | **t(df) = 2.72 (9,140)** | **< 0.01** |
| **Pubertal Timing (BL)**  Median (Q1-Q3) | -0.51 (1.59) | -0.14 (3.65) | **z = 3.53** | **< 0.001** |
| **CBCL Withdrawn/Depressed (BL)** Median (Q1-Q3) | 0 (0-2) | 0 (0-1) | **z = 5.54** | **< 0.001** |
| **CBCL Withdrawn/Depressed (Y2)** Median (Q1-Q3) | 0 (0-2) | 0 (0-2) | z = 0.20 | 0.84 |
| **CBCL Withdrawn/Depressed (Y3)** Median (Q1-Q3) | 1 (0-2) | 1 (0-2) | **z = −2.87** | **< 0.01** |

Lilliefors tests indicated non-normal distributions for pubertal timing (males: skewness = 1.43, kurtosis = 4.99; females: skewness = 0.30, kurtosis = 2.60) and for CBCL at all waves (all p < 0.001). Levene’s tests indicated unequal variances for pubertal timing and CBCL (all p < 0.001). BL: Baseline; Y2: 2-year follow-up; Y3: 3-year follow-up

#### **Table S2. Cross-sectional associations between pubertal timing and rsFC at baseline**

| **Predictor** | **Pooled** | | **Females** | | **Males** | |
| --- | --- | --- | --- | --- | --- | --- |
|  | **β** | **p** | **β** | **p** | **β** | **p** |
| Pubertal timing | -0.001 | 0.92 | 0.004 | 0.56 | -0.001 | 0.89 |
| Pubertal timing × Sex | 0.006 | 0.53 | – | – | – | – |
| Sex | **-0.020** | **0.01** | – | – | – | – |
| Age | **-0.012** | **<0.01** | -0.011 | 0.09 | **-0.011** | **0.02** |
| Network-pair slope SD | 0.033 | | 0.042 | | 0.021 | |

#### **Table S3. Model comparison of shared and sex-specific variability in network-pair–specific baseline pubertal timing–rsFC associations**

| **Model** | **Specification** | **Variance Components (SD)** | **AIC** | **BIC** | **Likelihood Ratio Test** |
| --- | --- | --- | --- | --- | --- |
| **Pooled model** | Network-pair–specific slopes (shared across sexes) | 0.033 | 3,535,438 | 3,535,559 | **χ²(1) = 40.32**  **p < .001** |
| **Extended model** | Adds female-specific deviations in network-pair slopes | Male: 0.029  Female deviation: 0.020 🡪 total Female ≈ 0.035 | 3,535,400 | 3,535,533 |  |

AIC: Akaike Information Criterion; BIC: Bayesian Information Criterion;

#### **Table S4. Network-pair–specific associations between pubertal timing and resting-state functional connectivity at baseline in females**

| **Network Pairs** | **β (median)** | **95% CrI lower** | **95% CrI upper** |
| --- | --- | --- | --- |
| AMYG_AMYG | -0.011 | -0.041 | 0.020 |
| AMYG_BG | -0.019 | -0.049 | 0.011 |
| **AMYG_CEREB** | **0.032** | **0.003** | **0.063** |
| AMYG_THAL | -0.015 | -0.046 | 0.014 |
| AUD_AMYG | 0.021 | -0.008 | 0.051 |
| AUD_AUD | -0.004 | -0.035 | 0.027 |
| **AUD_BG** | **-0.090** | **-0.120** | **-0.060** |
| **AUD_CAN** | **0.041** | **0.011** | **0.072** |
| AUD_CEREB | -0.028 | -0.057 | 0.002 |
| AUD_HIPP | 0.024 | -0.007 | 0.053 |
| **AUD_PMN** | **-0.033** | **-0.064** | **-0.001** |
| AUD_THAL | -0.025 | -0.056 | 0.005 |
| BG_BG | 0.002 | -0.029 | 0.032 |
| **BG_CEREB** | **0.080** | **0.050** | **0.110** |
| **BG_THAL** | **0.036** | **0.006** | **0.066** |
| CAN_AMYG | -0.027 | -0.058 | 0.002 |
| CAN_BG | 0.010 | -0.021 | 0.042 |
| **CAN_CAN** | **-0.052** | **-0.081** | **-0.021** |
| CAN_CEREB | 0.016 | -0.014 | 0.048 |
| CAN_HIPP | -0.016 | -0.047 | 0.013 |
| CAN_THAL | -0.007 | -0.037 | 0.023 |
| **CEREB_CEREB** | **0.037** | **0.007** | **0.067** |
| CON_AMYG | 0.031 | 0.000 | 0.062 |
| **CON_AUD** | **-0.050** | **-0.081** | **-0.019** |
| CON_BG | -0.030 | -0.060 | 0.002 |
| **CON_CAN** | **0.053** | **0.023** | **0.085** |
| **CON_CEREB** | **0.041** | **0.009** | **0.072** |
| **CON_CON** | **-0.077** | **-0.108** | **-0.047** |
| CON_HIPP | 0.024 | -0.007 | 0.054 |
| CON_PMN | 0.009 | -0.022 | 0.040 |
| **CON_SMD** | **-0.031** | **-0.063** | **-0.001** |
| CON_SML | -0.024 | -0.055 | 0.006 |
| CON_THAL | 0.001 | -0.030 | 0.030 |
| DAN_AMYG | 0.016 | -0.016 | 0.047 |
| **DAN_AUD** | **0.037** | **0.007** | **0.067** |
| **DAN_BG** | **0.036** | **0.005** | **0.067** |
| DAN_CAN | 0.009 | -0.023 | 0.040 |
| **DAN_CEREB** | **0.039** | **0.008** | **0.070** |
| DAN_CON | 0.000 | -0.030 | 0.031 |
| **DAN_DAN** | **-0.037** | **-0.067** | **-0.004** |
| **DAN_HIPP** | **0.055** | **0.024** | **0.086** |
| DAN_PMN | 0.000 | -0.030 | 0.031 |
| **DAN_SAL** | **0.040** | **0.009** | **0.071** |
| DAN_SMD | 0.001 | -0.030 | 0.033 |
| DAN_SML | 0.023 | -0.008 | 0.054 |
| DAN_THAL | 0.002 | -0.030 | 0.034 |
| **DAN_VAN** | **0.060** | **0.031** | **0.090** |
| DMN_AMYG | -0.024 | -0.054 | 0.007 |
| **DMN_AUD** | **-0.033** | **-0.066** | **-0.002** |
| DMN_BG | 0.003 | -0.028 | 0.035 |
| DMN_CAN | -0.015 | -0.047 | 0.016 |
| **DMN_CEREB** | **0.046** | **0.015** | **0.077** |
| **DMN_CON** | **0.037** | **0.007** | **0.067** |
| **DMN_DAN** | **0.059** | **0.027** | **0.090** |
| **DMN_DMN** | **-0.030** | **-0.061** | **-0.001** |
| **DMN_FPN** | **0.037** | **0.006** | **0.068** |
| **DMN_HIPP** | **-0.035** | **-0.065** | **-0.004** |
| DMN_PMN | 0.005 | -0.025 | 0.036 |
| DMN_SAL | -0.007 | -0.036 | 0.023 |
| DMN_SCAN | -0.022 | -0.054 | 0.008 |
| DMN_SMD | -0.020 | -0.051 | 0.009 |
| **DMN_SML** | **-0.033** | **-0.063** | **-0.002** |
| DMN_THAL | 0.028 | -0.003 | 0.059 |
| **DMN_VAN** | **-0.049** | **-0.080** | **-0.019** |
| DMN_VIS | 0.022 | -0.008 | 0.052 |
| FPN_AMYG | 0.000 | -0.030 | 0.031 |
| **FPN_AUD** | **-0.036** | **-0.067** | **-0.004** |
| **FPN_BG** | **0.047** | **0.016** | **0.077** |
| **FPN_CAN** | **0.034** | **0.004** | **0.065** |
| **FPN_CEREB** | **0.059** | **0.028** | **0.089** |
| FPN_CON | 0.025 | -0.006 | 0.057 |
| FPN_DAN | -0.013 | -0.043 | 0.017 |
| FPN_FPN | 0.028 | -0.003 | 0.060 |
| FPN_HIPP | 0.023 | -0.008 | 0.053 |
| **FPN_PMN** | **0.040** | **0.009** | **0.070** |
| **FPN_SAL** | **0.041** | **0.011** | **0.072** |
| **FPN_SMD** | **-0.049** | **-0.079** | **-0.019** |
| **FPN_SML** | **-0.036** | **-0.067** | **-0.005** |
| **FPN_THAL** | **0.052** | **0.020** | **0.082** |
| FPN_VAN | 0.008 | -0.023 | 0.038 |
| HIPP_AMYG | -0.010 | -0.039 | 0.021 |
| HIPP_BG | -0.021 | -0.053 | 0.009 |
| HIPP_CEREB | -0.017 | -0.048 | 0.014 |
| HIPP_HIPP | -0.021 | -0.050 | 0.009 |
| HIPP_THAL | 0.006 | -0.024 | 0.038 |
| PMN_AMYG | -0.022 | -0.053 | 0.008 |
| **PMN_BG** | **0.080** | **0.049** | **0.109** |
| PMN_CAN | -0.012 | -0.041 | 0.018 |
| **PMN_CEREB** | **0.090** | **0.059** | **0.122** |
| PMN_HIPP | -0.026 | -0.058 | 0.006 |
| PMN_PMN | 0.001 | -0.028 | 0.032 |
| **PMN_THAL** | **0.056** | **0.026** | **0.087** |
| SAL_AMYG | -0.005 | -0.036 | 0.025 |
| **SAL_AUD** | **-0.058** | **-0.088** | **-0.028** |
| SAL_BG | -0.015 | -0.045 | 0.015 |
| **SAL_CAN** | **0.034** | **0.004** | **0.065** |
| **SAL_CEREB** | **0.079** | **0.048** | **0.111** |
| SAL_CON | -0.016 | -0.047 | 0.015 |
| **SAL_HIPP** | **-0.032** | **-0.061** | **0.000** |
| **SAL_PMN** | **0.047** | **0.016** | **0.076** |
| **SAL_SAL** | **-0.033** | **-0.064** | **-0.002** |
| **SAL_SMD** | **-0.054** | **-0.084** | **-0.023** |
| **SAL_SML** | **-0.048** | **-0.077** | **-0.016** |
| **SAL_THAL** | **0.034** | **0.005** | **0.065** |
| **SCAN_AMYG** | **0.041** | **0.011** | **0.073** |
| **SCAN_AUD** | **0.053** | **0.020** | **0.084** |
| **SCAN_BG** | **-0.048** | **-0.079** | **-0.017** |
| **SCAN_CAN** | **0.035** | **0.004** | **0.066** |
| SCAN_CEREB | -0.022 | -0.053 | 0.010 |
| **SCAN_CON** | **-0.034** | **-0.065** | **-0.004** |
| SCAN_DAN | -0.007 | -0.041 | 0.024 |
| **SCAN_FPN** | **-0.047** | **-0.077** | **-0.016** |
| **SCAN_HIPP** | **0.063** | **0.032** | **0.094** |
| **SCAN_PMN** | **-0.043** | **-0.075** | **-0.011** |
| **SCAN_SAL** | **-0.044** | **-0.073** | **-0.015** |
| SCAN_SCAN | 0.005 | -0.026 | 0.035 |
| **SCAN_SMD** | **0.110** | **0.080** | **0.141** |
| **SCAN_SML** | **0.030** | **0.000** | **0.060** |
| SCAN_THAL | -0.014 | -0.044 | 0.016 |
| SCAN_VAN | 0.009 | -0.022 | 0.040 |
| SCAN_VIS | 0.021 | -0.010 | 0.051 |
| SMD_AMYG | 0.028 | -0.003 | 0.059 |
| **SMD_AUD** | **0.059** | **0.028** | **0.089** |
| **SMD_BG** | **-0.060** | **-0.090** | **-0.031** |
| **SMD_CAN** | **0.054** | **0.023** | **0.086** |
| **SMD_CEREB** | **-0.068** | **-0.099** | **-0.037** |
| **SMD_HIPP** | **0.038** | **0.009** | **0.069** |
| SMD_PMN | -0.031 | -0.063 | 0.000 |
| **SMD_SMD** | **0.066** | **0.035** | **0.097** |
| **SMD_SML** | **0.103** | **0.073** | **0.134** |
| SMD_THAL | -0.010 | -0.041 | 0.021 |
| SML_AMYG | 0.018 | -0.013 | 0.049 |
| **SML_AUD** | **0.043** | **0.010** | **0.075** |
| **SML_BG** | **-0.079** | **-0.111** | **-0.048** |
| SML_CAN | 0.026 | -0.005 | 0.056 |
| **SML_CEREB** | **-0.047** | **-0.079** | **-0.017** |
| SML_HIPP | 0.028 | -0.004 | 0.058 |
| **SML_PMN** | **-0.043** | **-0.074** | **-0.013** |
| SML_SML | -0.004 | -0.035 | 0.026 |
| **SML_THAL** | **-0.059** | **-0.089** | **-0.028** |
| **THAL_CEREB** | **0.071** | **0.041** | **0.102** |
| **THAL_THAL** | **0.052** | **0.021** | **0.082** |
| VAN_AMYG | -0.005 | -0.035 | 0.025 |
| VAN_AUD | -0.003 | -0.032 | 0.027 |
| **VAN_BG** | **-0.077** | **-0.107** | **-0.046** |
| VAN_CAN | -0.001 | -0.031 | 0.029 |
| VAN_CEREB | 0.005 | -0.026 | 0.036 |
| VAN_CON | 0.008 | -0.023 | 0.040 |
| VAN_HIPP | -0.016 | -0.048 | 0.015 |
| VAN_PMN | -0.020 | -0.051 | 0.010 |
| **VAN_SAL** | **-0.069** | **-0.101** | **-0.038** |
| VAN_SMD | 0.011 | -0.020 | 0.043 |
| VAN_SML | 0.004 | -0.027 | 0.035 |
| VAN_THAL | -0.025 | -0.054 | 0.006 |
| VAN_VAN | -0.021 | -0.053 | 0.008 |
| VIS_AMYG | 0.028 | -0.003 | 0.058 |
| **VIS_AUD** | **0.086** | **0.055** | **0.118** |
| VIS_BG | 0.016 | -0.014 | 0.047 |
| **VIS_CAN** | **-0.031** | **-0.062** | **0.000** |
| VIS_CEREB | -0.002 | -0.033 | 0.028 |
| VIS_CON | 0.013 | -0.017 | 0.044 |
| VIS_DAN | -0.031 | -0.060 | 0.00006 |
| VIS_FPN | 0.007 | -0.023 | 0.036 |
| **VIS_HIPP** | **0.039** | **0.010** | **0.069** |
| VIS_PMN | -0.015 | -0.045 | 0.015 |
| VIS_SAL | -0.002 | -0.033 | 0.030 |
| **VIS_SMD** | **0.079** | **0.048** | **0.110** |
| **VIS_SML** | **0.051** | **0.019** | **0.082** |
| **VIS_THAL** | **-0.040** | **-0.071** | **-0.008** |
| **VIS_VAN** | **0.075** | **0.045** | **0.107** |
| **VIS_VIS** | **-0.059** | **-0.089** | **-0.028** |

#### **Table S5. Network-pair–specific associations between pubertal timing and resting-state functional connectivity at baseline in males**

| **Network Pairs** | | **β (median)** | | **95% CrI lower** | | **95% CrI upper** |
| --- | --- | --- | --- | --- | --- | --- |
| AMYG_AMYG | 0.003 | | -0.022 | | 0.028 | |
| AMYG_BG | -0.007 | | -0.033 | | 0.019 | |
| AMYG_CEREB | 0.008 | | -0.018 | | 0.034 | |
| AMYG_THAL | 0.005 | | -0.020 | | 0.031 | |
| AUD_AMYG | 0.005 | | -0.020 | | 0.030 | |
| AUD_AUD | -0.002 | | -0.027 | | 0.024 | |
| AUD_BG | -0.018 | | -0.045 | | 0.008 | |
| **AUD_CAN** | **0.034** | | **0.008** | | **0.059** | |
| **AUD_CEREB** | **-0.035** | | **-0.061** | | **-0.008** | |
| AUD_HIPP | -0.019 | | -0.045 | | 0.007 | |
| AUD_PMN | -0.014 | | -0.040 | | 0.012 | |
| AUD_THAL | -0.011 | | -0.037 | | 0.015 | |
| BG_BG | 0.010 | | -0.016 | | 0.036 | |
| BG_CEREB | 0.001 | | -0.025 | | 0.028 | |
| BG_THAL | 0.001 | | -0.024 | | 0.027 | |
| CAN_AMYG | 0.000 | | -0.026 | | 0.025 | |
| CAN_BG | 0.010 | | -0.017 | | 0.037 | |
| **CAN_CAN** | **-0.031** | | **-0.056** | | **-0.006** | |
| CAN_CEREB | -0.004 | | -0.029 | | 0.022 | |
| CAN_HIPP | -0.001 | | -0.027 | | 0.024 | |
| CAN_THAL | -0.010 | | -0.037 | | 0.017 | |
| **CEREB_CEREB** | **0.034** | | **0.008** | | **0.061** | |
| CON_AMYG | -0.006 | | -0.032 | | 0.019 | |
| CON_AUD | -0.013 | | -0.038 | | 0.012 | |
| CON_BG | -0.018 | | -0.042 | | 0.008 | |
| **CON_CAN** | **0.035** | | **0.008** | | **0.061** | |
| CON_CEREB | -0.017 | | -0.042 | | 0.010 | |
| CON_CON | -0.021 | | -0.048 | | 0.005 | |
| CON_HIPP | -0.002 | | -0.026 | | 0.023 | |
| CON_PMN | 0.000 | | -0.027 | | 0.025 | |
| CON_SMD | -0.004 | | -0.030 | | 0.021 | |
| CON_SML | 0.001 | | -0.025 | | 0.027 | |
| CON_THAL | 0.003 | | -0.022 | | 0.029 | |
| DAN_AMYG | -0.008 | | -0.033 | | 0.017 | |
| DAN_AUD | 0.009 | | -0.018 | | 0.034 | |
| DAN_BG | 0.014 | | -0.011 | | 0.039 | |
| DAN_CAN | -0.018 | | -0.044 | | 0.009 | |
| DAN_CEREB | -0.010 | | -0.034 | | 0.016 | |
| DAN_CON | 0.013 | | -0.012 | | 0.040 | |
| DAN_DAN | -0.015 | | -0.041 | | 0.010 | |
| DAN_HIPP | 0.022 | | -0.003 | | 0.047 | |
| DAN_PMN | -0.001 | | -0.027 | | 0.024 | |
| **DAN_SAL** | **0.033** | | **0.006** | | **0.060** | |
| DAN_SMD | -0.006 | | -0.033 | | 0.019 | |
| DAN_SML | 0.014 | | -0.011 | | 0.041 | |
| DAN_THAL | -0.002 | | -0.027 | | 0.024 | |
| DAN_VAN | 0.019 | | -0.006 | | 0.044 | |
| DMN_AMYG | -0.010 | | -0.035 | | 0.016 | |
| DMN_AUD | -0.015 | | -0.041 | | 0.010 | |
| DMN_BG | -0.006 | | -0.032 | | 0.020 | |
| DMN_CAN | -0.011 | | -0.037 | | 0.014 | |
| DMN_CEREB | -0.009 | | -0.036 | | 0.017 | |
| DMN_CON | 0.009 | | -0.018 | | 0.035 | |
| **DMN_DAN** | **0.032** | | **0.007** | | **0.059** | |
| **DMN_DMN** | **-0.029** | | **-0.054** | | **-0.003** | |
| DMN_FPN | 0.021 | | -0.005 | | 0.046 | |
| DMN_HIPP | -0.015 | | -0.039 | | 0.010 | |
| DMN_PMN | 0.011 | | -0.014 | | 0.037 | |
| **DMN_SAL** | **-0.028** | | **-0.055** | | **-0.002** | |
| DMN_SCAN | -0.016 | | -0.041 | | 0.010 | |
| DMN_SMD | -0.001 | | -0.026 | | 0.025 | |
| DMN_SML | -0.009 | | -0.034 | | 0.017 | |
| DMN_THAL | -0.024 | | -0.050 | | 0.002 | |
| DMN_VAN | -0.013 | | -0.038 | | 0.012 | |
| DMN_VIS | 0.017 | | -0.010 | | 0.043 | |
| FPN_AMYG | -0.006 | | -0.031 | | 0.020 | |
| **FPN_AUD** | **-0.031** | | **-0.057** | | **-0.006** | |
| FPN_BG | 0.003 | | -0.022 | | 0.028 | |
| FPN_CAN | 0.005 | | -0.020 | | 0.030 | |
| FPN_CEREB | -0.001 | | -0.027 | | 0.026 | |
| FPN_CON | -0.003 | | -0.028 | | 0.022 | |
| FPN_DAN | 0.004 | | -0.021 | | 0.029 | |
| FPN_FPN | 0.001 | | -0.023 | | 0.026 | |
| FPN_HIPP | -0.009 | | -0.036 | | 0.017 | |
| FPN_PMN | 0.001 | | -0.024 | | 0.027 | |
| FPN_SAL | -0.013 | | -0.039 | | 0.012 | |
| FPN_SMD | -0.019 | | -0.046 | | 0.007 | |
| FPN_SML | -0.022 | | -0.046 | | 0.003 | |
| FPN_THAL | -0.019 | | -0.043 | | 0.007 | |
| FPN_VAN | 0.012 | | -0.013 | | 0.038 | |
| HIPP_AMYG | -0.009 | | -0.035 | | 0.018 | |
| HIPP_BG | -0.008 | | -0.034 | | 0.018 | |
| HIPP_CEREB | -0.009 | | -0.035 | | 0.016 | |
| HIPP_HIPP | 0.000 | | -0.026 | | 0.026 | |
| HIPP_THAL | -0.005 | | -0.032 | | 0.022 | |
| PMN_AMYG | -0.012 | | -0.038 | | 0.013 | |
| PMN_BG | 0.015 | | -0.011 | | 0.041 | |
| PMN_CAN | -0.009 | | -0.034 | | 0.015 | |
| PMN_CEREB | 0.010 | | -0.016 | | 0.036 | |
| PMN_HIPP | -0.001 | | -0.026 | | 0.024 | |
| PMN_PMN | -0.012 | | -0.038 | | 0.013 | |
| PMN_THAL | 0.008 | | -0.017 | | 0.035 | |
| SAL_AMYG | -0.023 | | -0.049 | | 0.002 | |
| SAL_AUD | -0.005 | | -0.031 | | 0.020 | |
| SAL_BG | 0.008 | | -0.018 | | 0.033 | |
| SAL_CAN | 0.017 | | -0.009 | | 0.043 | |
| SAL_CEREB | -0.004 | | -0.029 | | 0.021 | |
| SAL_CON | 0.010 | | -0.016 | | 0.036 | |
| SAL_HIPP | -0.016 | | -0.041 | | 0.010 | |
| SAL_PMN | -0.009 | | -0.035 | | 0.015 | |
| SAL_SAL | -0.023 | | -0.047 | | 0.003 | |
| SAL_SMD | 0.005 | | -0.021 | | 0.031 | |
| SAL_SML | 0.003 | | -0.023 | | 0.029 | |
| SAL_THAL | -0.005 | | -0.030 | | 0.020 | |
| SCAN_AMYG | -0.004 | | -0.029 | | 0.022 | |
| **SCAN_AUD** | **0.033** | | **0.008** | | **0.059** | |
| SCAN_BG | -0.013 | | -0.038 | | 0.012 | |
| SCAN_CAN | 0.019 | | -0.006 | | 0.044 | |
| SCAN_CEREB | -0.020 | | -0.045 | | 0.006 | |
| SCAN_CON | 0.016 | | -0.010 | | 0.041 | |
| SCAN_DAN | 0.004 | | -0.022 | | 0.029 | |
| **SCAN_FPN** | **-0.026** | | **-0.053** | | **0.000** | |
| SCAN_HIPP | 0.020 | | -0.005 | | 0.046 | |
| SCAN_PMN | -0.009 | | -0.036 | | 0.017 | |
| SCAN_SAL | 0.013 | | -0.013 | | 0.040 | |
| SCAN_SCAN | 0.017 | | -0.008 | | 0.043 | |
| **SCAN_SMD** | **0.040** | | **0.015** | | **0.066** | |
| SCAN_SML | 0.019 | | -0.007 | | 0.046 | |
| SCAN_THAL | 0.020 | | -0.006 | | 0.045 | |
| SCAN_VAN | -0.015 | | -0.040 | | 0.010 | |
| SCAN_VIS | -0.011 | | -0.037 | | 0.016 | |
| SMD_AMYG | 0.007 | | -0.019 | | 0.034 | |
| SMD_AUD | 0.018 | | -0.007 | | 0.045 | |
| SMD_BG | -0.015 | | -0.041 | | 0.010 | |
| **SMD_CAN** | **0.035** | | **0.011** | | **0.061** | |
| **SMD_CEREB** | **-0.031** | | **-0.056** | | **-0.005** | |
| SMD_HIPP | 0.000 | | -0.026 | | 0.026 | |
| SMD_PMN | 0.001 | | -0.025 | | 0.028 | |
| SMD_SMD | 0.018 | | -0.009 | | 0.044 | |
| **SMD_SML** | **0.035** | | **0.009** | | **0.061** | |
| SMD_THAL | 0.010 | | -0.016 | | 0.036 | |
| SML_AMYG | 0.013 | | -0.013 | | 0.039 | |
| SML_AUD | 0.023 | | -0.002 | | 0.049 | |
| SML_BG | -0.013 | | -0.038 | | 0.013 | |
| SML_CAN | 0.021 | | -0.004 | | 0.048 | |
| **SML_CEREB** | **-0.044** | | **-0.071** | | **-0.018** | |
| SML_HIPP | 0.003 | | -0.024 | | 0.028 | |
| SML_PMN | -0.012 | | -0.038 | | 0.014 | |
| SML_SML | 0.003 | | -0.022 | | 0.030 | |
| SML_THAL | -0.008 | | -0.034 | | 0.018 | |
| THAL_CEREB | 0.010 | | -0.016 | | 0.037 | |
| **THAL_THAL** | **0.035** | | **0.009** | | **0.061** | |
| VAN_AMYG | -0.006 | | -0.032 | | 0.019 | |
| VAN_AUD | -0.009 | | -0.036 | | 0.016 | |
| VAN_BG | -0.022 | | -0.048 | | 0.004 | |
| VAN_CAN | 0.017 | | -0.009 | | 0.043 | |
| **VAN_CEREB** | **-0.027** | | **-0.054** | | **0.000** | |
| VAN_CON | -0.015 | | -0.041 | | 0.013 | |
| VAN_HIPP | 0.002 | | -0.023 | | 0.028 | |
| VAN_PMN | 0.005 | | -0.021 | | 0.031 | |
| VAN_SAL | -0.016 | | -0.043 | | 0.010 | |
| VAN_SMD | -0.007 | | -0.032 | | 0.019 | |
| VAN_SML | -0.012 | | -0.037 | | 0.014 | |
| VAN_THAL | -0.022 | | -0.048 | | 0.004 | |
| VAN_VAN | -0.006 | | -0.031 | | 0.020 | |
| VIS_AMYG | 0.021 | | -0.005 | | 0.046 | |
| **VIS_AUD** | **0.036** | | **0.011** | | **0.062** | |
| VIS_BG | 0.015 | | -0.010 | | 0.041 | |
| VIS_CAN | -0.008 | | -0.034 | | 0.016 | |
| VIS_CEREB | 0.009 | | -0.017 | | 0.035 | |
| VIS_CON | 0.009 | | -0.018 | | 0.036 | |
| VIS_DAN | -0.018 | | -0.044 | | 0.007 | |
| VIS_FPN | -0.006 | | -0.032 | | 0.019 | |
| **VIS_HIPP** | **0.032** | | **0.006** | | **0.058** | |
| VIS_PMN | -0.012 | | -0.038 | | 0.013 | |
| VIS_SAL | 0.008 | | -0.018 | | 0.033 | |
| VIS_SMD | 0.019 | | -0.007 | | 0.046 | |
| VIS_SML | 0.014 | | -0.011 | | 0.039 | |
| VIS_THAL | 0.014 | | -0.011 | | 0.042 | |
| VIS_VAN | 0.017 | | -0.009 | | 0.043 | |
| **VIS_VIS** | **-0.037** | | **-0.062** | | **-0.010** | |

#### **Table S6. Cross-sectional and longitudinal associations between pubertal timing and depressive symptoms**

| **Predictor** | **CBCL Withdrawn/Depressed (BL)** | | **CBCL Withdrawn/Depressed (Y3)** | |
| --- | --- | --- | --- | --- |
|  | **β** | **p** | **β** | **p** |
| Pubertal timing | **0.17** | **<0.001** | **0.10** | **<0.01** |
| Sex | **-0.19** | **<0.001** | **0.21** | **<0.001** |
| Pubertal timing × Sex | -0.02 | 0.56 | -0.01 | 0.85 |
| Age | **0.04** | **0.02** | 0.01 | 0.15 |
| CBCL Withdrawn/Depressed (Y2) | – | – | **0.70** | **<0.001** |

BL: Baseline; Y2: 2-year follow-up; Y3: 3-year follow-up

#### **Table S7. Longitudinal associations between pubertal timing and rsFC**

| **Predictor** | **Females** | | **Males** | |
| --- | --- | --- | --- | --- |
|  | **β** | **p** | **β** | **p** |
| Pubertal timing | **-0.008** | **0.02** | -0.005 | 0.10 |
| rsFC_Y2_ | **0.277** | **<0.001** | **0.264** | **<0.001** |
| Age | 0.001 | 0.81 | -0.006 | 0.06 |
| Network-pair slope SD | 0.036 | | ~0 | |

Y2: 2-year follow-up

**Table S8. Fold-wise stability of network-pair–specific associations between pubertal timing and rsFC at the 2-year follow-up in females.** Only network pairs that were selected in at least one cross-validation fold are shown.

| **Network Pairs** | **Folds selected (of 5)** | **Direction (stable)** |
| --- | --- | --- |
| AMYG_AMYG | 1 | - |
| AMYG_BG | 2 | - |
| **AMYG_THAL** | **3** | **Negative** |
| **AUD_BG** | **5** | **Negative** |
| **AUD_CEREB** | **3** | **Negative** |
| AUD_HIPP | 2 | - |
| **AUD_THAL** | **4** | **Negative** |
| **BG_CEREB** | **5** | **Positive** |
| **CON_AUD** | **4** | **Negative** |
| CON_CAN | 2 | - |
| CON_CON | 1 | - |
| **CON_SMD** | **4** | **Negative** |
| **CON_SML** | **3** | **Negative** |
| DAN_HIPP | 2 | - |
| DMN_SAL | 1 | - |
| FPN_SMD | 1 | - |
| FPN_SML | 1 | - |
| **FPN_VAN** | **3** | **Negative** |
| **HIPP_AMYG** | **4** | **Negative** |
| HIPP_BG | 1 | - |
| HIPP_CEREB | 1 | - |
| **PMN_BG** | **3** | **Positive** |
| PMN_CEREB | 1 | - |
| **PMN_THAL** | **4** | **Positive** |
| **SAL_AUD** | **4** | **Negative** |
| SAL_CAN | 1 | - |
| SAL_HIPP | 1 | - |
| SAL_PMN | 1 | - |
| **SAL_SML** | **5** | **Negative** |
| SCAN_BG | 2 | - |
| SCAN_CEREB | 1 | - |
| **SMD_BG** | **5** | **Negative** |
| **SMD_CEREB** | **3** | **Negative** |
| SMD_SML | 2 | - |
| **SMD_THAL** | **5** | **Negative** |
| **SML_BG** | **5** | **Negative** |
| **SML_CEREB** | **5** | **Negative** |
| SML_THAL | 2 | - |
| **THAL_CEREB** | **3** | **Positive** |
| **VAN_AUD** | **5** | **Negative** |
| **VAN_BG** | **5** | **Negative** |
| **VAN_CON** | **5** | **Negative** |
| VAN_PMN | 1 | - |
| **VAN_SAL** | **4** | **Negative** |
| VAN_SML | 2 | - |
| **VAN_THAL** | **4** | **Negative** |
| Vis_AUD | 2 | - |
| Vis_SMD | 2 | - |
| Vis_VAN | 1 | - |

#### **Table S9. Network-pair–specific mediation effects linking pubertal timing, rsFC at the 2-year follow-up, and depressive symptoms at the 3-year follow-up in females**

| **Network Pairs** | **a** | **b** | **Indirect effect (a × b)** | **95% CrI lower** | **95% CrI upper** | **Direct effect (c′)** | **Total effect (c)** |
| --- | --- | --- | --- | --- | --- | --- | --- |
| AMYG_THAL | **-0.053** | 0.000058 | -0.000002 | -0.000746 | 0.000715 | **0.398** | **0.398350** |
| AUD_BG | **-0.090** | -0.000045 | 0.000004 | -0.001124 | 0.001190 | **0.398** | **0.398562** |
| AUD_CEREB | **-0.061** | 0.000076 | -0.000001 | -0.000831 | 0.000856 | **0.398** | **0.398338** |
| AUD_THAL | **-0.066** | -0.000081 | 0.000004 | -0.000864 | 0.000882 | **0.398** | **0.398313** |
| BG_CEREB | **0.085** | 0.000070 | 0.000005 | -0.001123 | 0.001082 | **0.398** | **0.397953** |
| CON_AUD | **-0.068** | 0.000198 | -0.000009 | -0.000902 | 0.000907 | **0.398** | **0.398498** |
| CON_SMD | **-0.060** | 0.000169 | -0.000007 | -0.000863 | 0.000829 | **0.398** | **0.398347** |
| CON_SML | **-0.060** | 0.000109 | -0.000004 | -0.000856 | 0.000784 | **0.398** | **0.398440** |
| FPN_VAN | **-0.051** | 0.000026 | 0.000001 | -0.000723 | 0.000752 | **0.398** | **0.398393** |
| HIPP_AMYG | **-0.059** | 0.000060 | -0.000003 | -0.000776 | 0.000759 | **0.398** | **0.398401** |
| PMN_BG | **0.067** | 0.000095 | 0.000004 | -0.000854 | 0.000878 | **0.398** | **0.398141** |
| PMN_THAL | **0.073** | 0.000214 | 0.000014 | -0.000980 | 0.000975 | **0.398** | **0.398143** |
| SAL_AUD | **-0.058** | 0.000150 | -0.000007 | -0.000870 | 0.000787 | **0.398** | **0.398523** |
| SAL_SML | **-0.092** | 0.000176 | -0.000015 | -0.001164 | 0.001154 | **0.398** | **0.398562** |
| SMD_BG | **-0.075** | 0.000023 | -0.000002 | -0.000995 | 0.000990 | **0.398** | **0.398374** |
| SMD_CEREB | **-0.060** | -0.000033 | 0.000002 | -0.000805 | 0.000817 | **0.398** | **0.398334** |
| SMD_THAL | **-0.076** | 0.000209 | -0.000013 | -0.000994 | 0.000986 | **0.398** | **0.398397** |
| SML_BG | **-0.095** | -0.000034 | 0.000004 | -0.001164 | 0.001240 | **0.398** | **0.398428** |
| SML_CEREB | **-0.072** | -0.000011 | 0.000001 | -0.000935 | 0.000980 | **0.398** | **0.398437** |
| THAL_CEREB | **0.064** | -0.000060 | -0.000003 | -0.000880 | 0.000823 | **0.398** | **0.398223** |
| VAN_AUD | **-0.067** | 0.000146 | -0.000009 | -0.000940 | 0.000859 | **0.398** | **0.398441** |
| VAN_BG | **-0.082** | 0.000069 | -0.000005 | -0.001105 | 0.001065 | **0.398** | **0.398194** |
| VAN_CON | **-0.067** | 0.000129 | -0.000007 | -0.000865 | 0.000923 | **0.398** | **0.398365** |
| VAN_SAL | **-0.064** | 0.000128 | -0.000006 | -0.000917 | 0.000855 | **0.398** | **0.398423** |
| VAN_THAL | **-0.061** | 0.000057 | -0.000003 | -0.000844 | 0.000839 | **0.398** | **0.398415** |

a: Pubertal timing 🡪 rsFC_Y2_; b: rsFC_Y2_ 🡪 depression_Y3_; Bold values denote estimates with 95% credible intervals that exclude zero.

### References

1.	Dosenbach NUF, Koller JM, Earl EA, Miranda-Dominguez O, Klein RL, Van AN, et al. Real-time motion analytics during brain MRI improve data quality and reduce costs [Internet]. 2017. doi:10.1016/j.neuroimage.2017.08.025

2.	Casey BJ, Cannonier T, Conley MI, Cohen AO, Barch DM, Heitzeg MM, et al. The Adolescent Brain Cognitive Development (ABCD) study: Imaging acquisition across 21 sites. Developmental Cognitive Neuroscience. Elsevier Ltd; 2018. p. 43–54. doi:10.1016/j.dcn.2018.03.001 PubMed PMID: 29567376.

3.	Feczko E, Conan G, Marek S, Tervo-Clemmens B, Cordova M, Doyle O, et al. Adolescent Brain Cognitive Development (ABCD) Community MRI Collection and Utilities. bioRxiv. 2021 Jul 11;20:2021.07.09.451638. doi:10.1101/2021.07.09.451638

4.	Power JD, Barnes KA, Snyder AZ, Schlaggar BL, Petersen SE. Spurious but systematic correlations in functional connectivity MRI networks arise from subject motion. Neuroimage. 2012;59:2142–54. doi:10.1016/j.neuroimage.2011.10.018

5.	Power JD, Mitra A, Laumann TO, Snyder AZ, Schlaggar BL, Petersen SE. Methods to detect, characterize, and remove motion artifact in resting state fMRI. Neuroimage. 2014 Jan 1;84:320–41. doi:10.1016/J.NEUROIMAGE.2013.08.048 PubMed PMID: 23994314.

6.	Hallquist MN, Hwang K, Luna B. The nuisance of nuisance regression: Spectral misspecification in a common approach to resting-state fMRI preprocessing reintroduces noise and obscures functional connectivity ☆ [Internet]. 2013. doi:10.1016/j.neuroimage.2013.05.116

7.	Carp J. Optimizing the order of operations for movement scrubbing: Comment on Power et al. Neuroimage. 2013;76:436–8. doi:10.1016/j.neuroimage.2011.12.061

8.	Gordon EM, Laumann TO, Adeyemo B, Huckins JF, Kelley WM, Petersen SE. Generation and Evaluation of a Cortical Area Parcellation from Resting-State Correlations. Cerebral Cortex. 2016 Jan 1;26(1):288–303. doi:10.1093/CERCOR/BHU239 PubMed PMID: 25316338.

9.	Seitzman BA, Gratton C, Marek S, Raut R V., Dosenbach NUF, Schlaggar BL, et al. A set of functionally-defined brain regions with improved representation of the subcortex and cerebellum. Neuroimage. 2020 Feb 1;206:116290. doi:10.1016/J.NEUROIMAGE.2019.116290 PubMed PMID: 31634545.

10.	Fisher RA. Frequency Distribution of the Values of the Correlation Coefficient in Samples from an Indefinitely Large Population. Biometrika. 1915 May;10(4):507. doi:10.2307/2331838

11.	The MathWorks Inc. MATLAB. Natick, Massachusetts: The MathWorks Inc.; 2023.

12.	Achenbach TM, Rescorla LA. Manual for the ASEBA preschool forms and profiles. Burlington, VT: University of Vermont, Research center for children, youth, & families. 2000.

13.	van Duijvenvoorde ACK, Westhoff B, de Vos F, Wierenga LM, Crone EA. A three-wave longitudinal study of subcortical–cortical resting-state connectivity in adolescence: Testing age- and puberty-related changes. Hum Brain Mapp. 2019;40(13):3769–83. doi:10.1002/hbm.24630 PubMed PMID: 31099959.

14.	Ernst M, Benson B, Artiges E, Gorka AX, Lemaitre H, Lago T, et al. Pubertal maturation and sex effects on the default-mode network connectivity implicated in mood dysregulation. Transl Psychiatry. 2019;103(9). doi:10.1038/s41398-019-0433-6

15.	Duffy KA, Wiglesworth A, Roediger DJ, Mueller BA, Luciana M, Klimes-Dougan B, et al. Characterizing the effects of age, puberty, and sex on variability in resting-state functional connectivity in late childhood and early adolescence. Neuroimage. 2025;(313):121238. doi:10.1016/j.neuroimage.2025.121238

16.	Gracia-Tabuenca Z, Moreno MB, Barrios FA, Alcauter S. Development of the brain functional connectome follows puberty-dependent nonlinear trajectories. Neuroimage. 2021;229:117769. doi:10.1016/j.neuroimage.2021.117769

17.	Bates D, Mächler M, Bolker BM, Walker SC. Fitting Linear Mixed-Effects Models Using lme4. J Stat Softw. 2015 Oct 7;67(1):1–48. doi:10.18637/JSS.V067.I01

18.	R Core Team. R: A language and environment for statistical computing. R Foundation for Statistical Computing, editor. Vienne, Austria. URL https://www.R-project.org/.; 2025.

19.	Akaike H. A New Look at the Statistical Model Identification. IEEE Trans Automat Contr. 1974;19(6):716–23. doi:10.1109/TAC.1974.1100705

20.	Schwarz G. Estimating the Dimension of a Model. Ann Statist. 1978 Mar 1;6(2):461–4. doi:10.1214/AOS/1176344136

21.	Bürkner PC. brms: An R Package for Bayesian Multilevel Models Using Stan. J Stat Softw. 2017 Aug 29;80:1–28. doi:10.18637/JSS.V080.I01

22.	Bürkner PC. Advanced Bayesian Multilevel Modeling with the R Package brms. R Journal. 2017 May 31;10(1):395–411. doi:10.32614/rj-2018-017

23.	Bürkner PC. Bayesian Item Response Modeling in R with brms and Stan. J Stat Softw. 2021 Nov 30;100(5):1–54. doi:10.18637/JSS.V100.I05

24.	Gelman A, Hill J, Yajima M. Why We (Usually) Don’t Have to Worry About Multiple Comparisons. J Res Educ Eff. 2012 Apr;5(2):189–211. doi:10.1080/19345747.2011.618213

25.	Kuhn M. Building Predictive Models in R Using the caret Package. J Stat Softw. 2008 Nov 10;28(5):1–26. doi:10.18637/JSS.V028.I05

26.	Gelman A, Jakulin A, Pittau MG, Su YS. A weakly informative default prior distribution for logistic and other regression models. Annals of Applied Statistics. 2008 Dec 1;2(4):1360–83. doi:10.1214/08-AOAS191

27.	Biro FM, Greenspan LC, Galvez MP, Pinney SM, Teitelbaum S, Windham GC, et al. Onset of breast development in a longitudinal cohort. Pediatrics. 2013;132(6):1019–27. doi:10.1542/peds.2012-3773 PubMed PMID: 24190685.

28.	Quek YH, Tam WWS, Zhang MWB, Ho RCM. Exploring the association between childhood and adolescent obesity and depression: a meta-analysis. Obesity Reviews. 2017 Jul 1;18(7):742–54. doi:10.1111/OBR.12535 PubMed PMID: 28401646.

29.	Maoz H, Goldstein T, Goldstein BI, Axelson DA, Fan J, Hickey MB, et al. The Effects of Parental Mood on Reports of Their Children’s Psychopathology. J Am Acad Child Adolesc Psychiatry [Internet]. 2014 [cited 2025 Aug 18];53(10):1111–22. Available from: www.jaacap.org
